# OsWHY1/OsTRXz/OsMORFs complex is essential for RNA modification and early chloroplast development in rice

**DOI:** 10.1101/2024.08.28.610128

**Authors:** Xiangzi Zheng, Qingzheng Lu, Yuling Luo, Jiaxuan Xu, Weiqi Wang, Min Tan, Dongmei Liao, Wuqiang Hong, Sirong Chen, Chuheng Lin, Xiaoli Wang, Chunlan Fan, Habiba, Xiaowei Wang, Yanyun Li, Yu Zhang, Wenfang Lin, Ying Miao

## Abstract

WHIRLY (WHY) proteins are single-stranded DNA/RNA-binding proteins that play multifaceted roles in various plant species. The regulatory mechanisms of WHY proteins in rice remains blank. Here we demonstrate that *OsWHY1* in rice is required for early chloroplast development. CRISPR/Cas9-generated *oswhy1* knockout lines displayed albino seedling phenotypes, abnormal chloroplast structure and comprised redox balance in leaves. OsWHY1 interacts with multiple plastid proteins, including the thioredoxin OsTRXz and two multiple organellar RNA editing factors (OsMORF8 and OsMORF9) in chloroplasts. Accordingly, several plastid genes dependent on plastid-encoded RNA polymerase (PEP) in the *oswhy1* mutants were significantly depressed at both transcript and protein levels. The editing of *rps14* transcripts and splicing of *rpl2,* along with their protein expression, were defective in the *oswhy1* mutants. OsWHY1 exhibited RNA-binding activity, specifically binding to *rps14* and *rpl2* precursor RNAs, which underscores its role as a post-transcriptional regulator essential for normal protein synthesis in chloroplasts. Loss-of- function mutants of either *OsWHY1* or *OsMORF9* and *OsTRXz* displayed albino phenotypes, disrupted H_2_O_2_ homeostasis, and defective RNA processing in *rps14* and *rpl2*, suggesting the OsWHY1-OsTRXz-OsMORFs regulatory module is vital for maintaining chloroplast stability and integrity through its RNA-binding activity and its role in recruiting OsTRXz and OsMORFs to ensure proper RNA modification.

**One sentence summary:** OsWHY1 is integral to chloroplast development in rice through its RNA-binding activity and its role in recruiting OsTRXz and OsMORFs to ensure proper RNA modification.

## Introduction

Chloroplast development is concertedly modulated by plastid and nuclear genes, with their transcriptional activities closely coordinated by anterograde and retrograde signaling pathways (Hernandez-Verdeja and Strand, 2018). RNA metabolism, including RNA editing, splicing, and stabilization, plays a critical role in post-transcriptional modification and chloroplast gene expression (Sun et al., 2016). These RNA processes are mediated by various chloroplast RNA- binding proteins encoded by nuclear genes. These RNA-binding proteins typically contain specific structural domains, such as RNA recognition motifs (RRM), pentatricopeptide repeat motifs (PPR), or chloroplast RNA splicing ribosome maturation domains (CRM), which facilitate RNA processing (Jacobs and Kuck, 2011). RRM is the most widespread RNA binding domain and is present in all chloroplast ribonucleoproteins as well as in some CRM or PPR proteins, which contribute to splicing, editing, and stability control of chloroplast RNAs (Ruwe et al., 2011; Wu et al., 2021). CRM proteins are the dominant players in chloroplast RNA splicing. PPR proteins form a huge family and play multiple functions in various RNA processes (Shikanai and Fujii, 2013).

In Arabidopsis and maize, numerous RNA-binding proteins involved in RNA splicing or editing have been identified (Jacobs and Kuck, 2011), but only a few have been characterized in rice. For example, several CRM domain-containing proteins, including OsCRS1, OsCFM2, OsCFM3, as well as OsCAF1, are involved in intron splicing of several chloroplast transcripts such as *atpF, rpl2, rps12, rps16, ndhA, ndhB*, and *ycf3*, thereby influencing chloroplast development (Asakura et al., 2008; Liu et al., 2016; Zhang et al., 2019; Zhang et al., 2020). Loss of function of these CRM proteins strongly results in the albino and lethality phenotype of the seedling. More than 20 plastid-localized rice PPR proteins have now been reported to be essential for early chloroplast development by regulating RNA editing or splicing events of many plastid transcripts (Meng et al., 2024). Their corresponding mutants exhibit varying degrees of abnormal chloroplast developmental phenotypes. For example, the OsPPR1, OsPPR4, OsPPR6, OsSLA4, OsSLC1, and WAL3 mutants exhibit a seedling-lethal albino phenotype (Gothandam et al., 2005; Asano et al., 2013; Tang et al., 2017; Wang et al., 2018; Lv et al., 2020; Lv et al., 2022), while the WSL, WSL4, WSL5, and YLWS mutants developed white-striped leaves before the five-leaf stage (Tan et al., 2014; Wang et al., 2017; Liu et al., 2018; Lan et al., 2023). In contrast, mutation of the PPR genes *OsPGL1* or *OsPGL12* resulted in a pale green leaf phenotype at all vegetative stages (Xiao et al., 2018a; Chen et al., 2019). However, the detailed regulatory functions of these RNA-binding proteins in rice remain nuclear.

WHIRLY1 (WHY1) is a small plant-specific single-stranded nucleotide binding protein (Prikryl et al., 2008; Miao et al., 2013) and mainly serves as a transcriptional factor regulating genes involved in defense, plant senescence or abiotic stresses such as chilling response (Desveaux et al., 2000; Desveaux et al., 2004; Miao et al., 2013; Zhuang et al., 2019). WHY1 can be detected in both plastids and nucleus by immunological methods (Grabowski et al., 2008; Ren et al., 2017), therefore WHY1 is proposed to be involved in retrograde signaling (Krause et al., 2012; Lin et al., 2024). In Arabidopsis, AtWHY1 is reported to interact with light-harvesting protein complex I (LHCA1) in chloroplasts and may facilitate the stability of photosystem I and light adaptation (Huang et al., 2017). AtWHY1 also interacts with chloroplast RNase H1 AtRNH1C, which works in concert with the recombinase RecA1 to promote DNA repair process and chloroplast genome integrity (Wang et al., 2021a). Knockout of AtWHY1 leads to redox stress and SA accumulation in mature or senescing leaves, but do not affect chloroplast development at the early stage of plant growth (Lin et al., 2019; Lin et al., 2020). In barely, knockdown of HvWHY1 mediated by RNAi delays chloroplast development by retarding assembly of plastid ribosomes and complexes of the photosynthetic apparatus at the early stages of primary foliage leaf development (Kucharewicz et al., 2017; Krupinska et al., 2019). Tomato SlWHY1 can bind to upstream of photosynthesis-related gene *SlPsbA* in chloroplast and starch metabolism-associated genes, SlAMY3-L and SlISA2, in nucleus to positively regulate resistance of plant to chilling stress (Zhuang et al., 2019; Zhuang et al., 2020). In contrast, ZmWHY1 in maize and OsWHY1 in rice have been documented to be essential for early chloroplast development (Prikryl et al., 2008; Qiu et al., 2022).

Notably, WHY1 also interacts with RNA and participates in intron splicing of some plastid genes, suggesting that WHY1 is a transcription factor with RNA-binding activity. Recent studies have highlighted the dual functionality of certain transcription factors (TFs), where they not only bind DNA to regulate gene transcription but also bind RNA, influencing post-transcriptional processes (Holmes et al., 2020; Oksuz et al., 2023). For instance, APIP5, a bZIP-type TF in rice, binds to DNA in the nucleus to repress the transcription of defense genes, while in the cytoplasm, it binds to mRNAs to modulate their stability and turnover, thus integrating transcriptional and post-transcriptional regulation in plant immunity (Zhang et al., 2022). To date, very few plant TFs have been reported to bind to mRNA to control gene expression. Mutation of *ZmWHY1* reduces *atpF* intron splicing and accumulation of plastid ribosomes (Prikryl et al., 2008), while loss of *OsWHY1* affects the splicing and editing of *ndhA* transcripts (Qiu et al., 2022), implying that they may play different functions on post-transcriptional processes of plastid genes. However, the regulatory mechanisms of OsWHY1 in regulating post-transcriptional modification of the plastid transcripts remain obscure.

In this study, we focus on OsWHY1, the rice orthologue of AtWHY1 and AtWHY3 (Krause et al., 2005). Using CRISPR/Cas9-generated *oswhy1* knockout lines, we demonstrate that *OsWHY1* is crucial for early chloroplast development. The GFP-fused nanotrap-mass spectrometry analysis and bimolecular fluorescence complementation (BiFC) assay detection show that OsWHY1 predominantly localizes in chloroplasts, where it interacts with key plastid proteins, including a plastid thioredoxin, OsTRXz, identified as a PEP-associated protein and multiple organellar RNA editing factors (OsMORF8 and OsMORF9). These interactions are essential for proper RNA editing and splicing of specific plastid genes, such as *rps14* and *rpl2*, which are necessary for chloroplast stability and function. Furthermore, by using REMSA assays, OsWHY1 exhibits RNA-binding activity, specifically binding to specific sites of *rps14* and *rpl2* pre-mRNAs, suggesting its role as a post-transcriptional regulator essential for normal protein synthesis in chloroplasts. Mutation of the *OsWHY1, OsMORF9,* or *OsTRXz* genes results in increased reactive oxygen species (ROS) accumulation, defective RNA processing, and significant albino seedling phenotypes, highlighting the importance of the OsWHY1/OsTRXz/OsMORFs regulatory module in maintaining chloroplast integrity.

## Results

### Phenotypic characterization of the *oswhy1* mutant

According to the phylogenetic analysis of Arabidopsis *(Arabidopsis thaliana)* WHIRLY (WHY) proteins against translated sequences of the plant EST databases, rice (*Oryza sativa* L. ssp. *japonica*) possesses two WHY proteins encoded by the two genes, LOC_Os06g05350 (*OsWHY1*) and LOC_Os02g06370 (*OsWHY2*), which are homologues of *AtWHY1, AtWHY3* and *AtWHY2*, respectively (Krause et al., 2005). The protein sequence of OsWHY1 contains a typical Whirly transcription factor motif (PF08536) analyzed by MOTIF Search (https://www.genome.jp/tools/motif/) and the highly conserved KGKAAL sequence (or KGKAAM in OsWHY1) as well as a potential autoregulatory domain (PAD) by comparison with the corresponding WHY1 amino acid sequences from Arabidopsis (AT1G14410), potato (AAF91282), maize (ACC60344) and barley (BAJ96655) (**Supplementary Figure S1A**). The bioinformatic software TargetP and iPSORT (Emanuelsson et al., 2007) predicted a plastid transit peptide (PTP) located at the N-terminal 50 amino acids (aa) of OsWHY1, while the program cNLS Mapper (Kosugi et al. 2009) found a putative nuclear localization signal (pNLS) at the N-terminal 31 amino acids of the Whirly domain of OsWHY1 (**Supplementary Figure S1B**). To investigate the subcellular localization of OsWHY1, full length OsWHY1 (OsWHY1 1-272aa) and its PTP deletion variant (OsWHY1 51-272aa) C-terminally fused with GFP were co-expressed with an RFP-tagged nuclear WRKY protein (OsWRKY44-RFP) (Habiba et al., 2021) in rice protoplasts by a transient transformation. Imaging by confocal microscope showed that the green fluorescence of OsWHY1 1-272aa-GFP appeared as speckles and was clearly co-localized with the autofluorescence of chlorophyll in the chloroplasts, but not distributed in the nucleus due to the failure of the GFP fluorescence to overlap with RFP signals (**Supplementary Figure S1C**). This indicates that full length OsWHY1 protein is predominantly targeted to the chloroplasts in rice protoplasts. By contrast, the N-terminal deletion form of OsWHY1 (OsWHY1 51-272aa-GFP) was strongly visualized in the nucleus, but not in chloroplasts, as presented by the overlap of the GFP and RFP fluorescence signals (**Supplementary Figure S1C**). These findings imply that OsWHY1’s chloroplast localization is reliant on its N-terminal PTP sequence, and The NLS sequence in the OsWHY1 mutant may drive OsWHY1 into the nucleus in the absence of PTP.

To genetically explore biological function of *OsWHY1* in rice development, CRISPR/Cas9-based gene editing technology was conducted to generate *OsWHY1* mutants under the background of *Oryza sativa. L. japonica* ZH11 and two independent transgenic *oswhy1* knockout lines were obtained. They were identified by genomic DNA sequencing and characterized by a 39bp deletion from position 27 to 65 or a 7bp deletion from position 69 to 75 in the targeted exon region (**Figure 1A)**, truncating the PTP-encoding sequence in the *OsWHY1* open reading frame or causing frameshift mutation at the target site. Both homozygous mutants existed an albino seedling phenotype and eventually died (**Figure 1B**). Consistent with the albino phenotype, total chlorophyll contents were dramatically decreased in the *oswhy1* lines compared to that of wild type (WT), which was accompanied by barely detectable photochemical efficiency in these mutants (**Figure 1C, D**).

**Figure 1.**
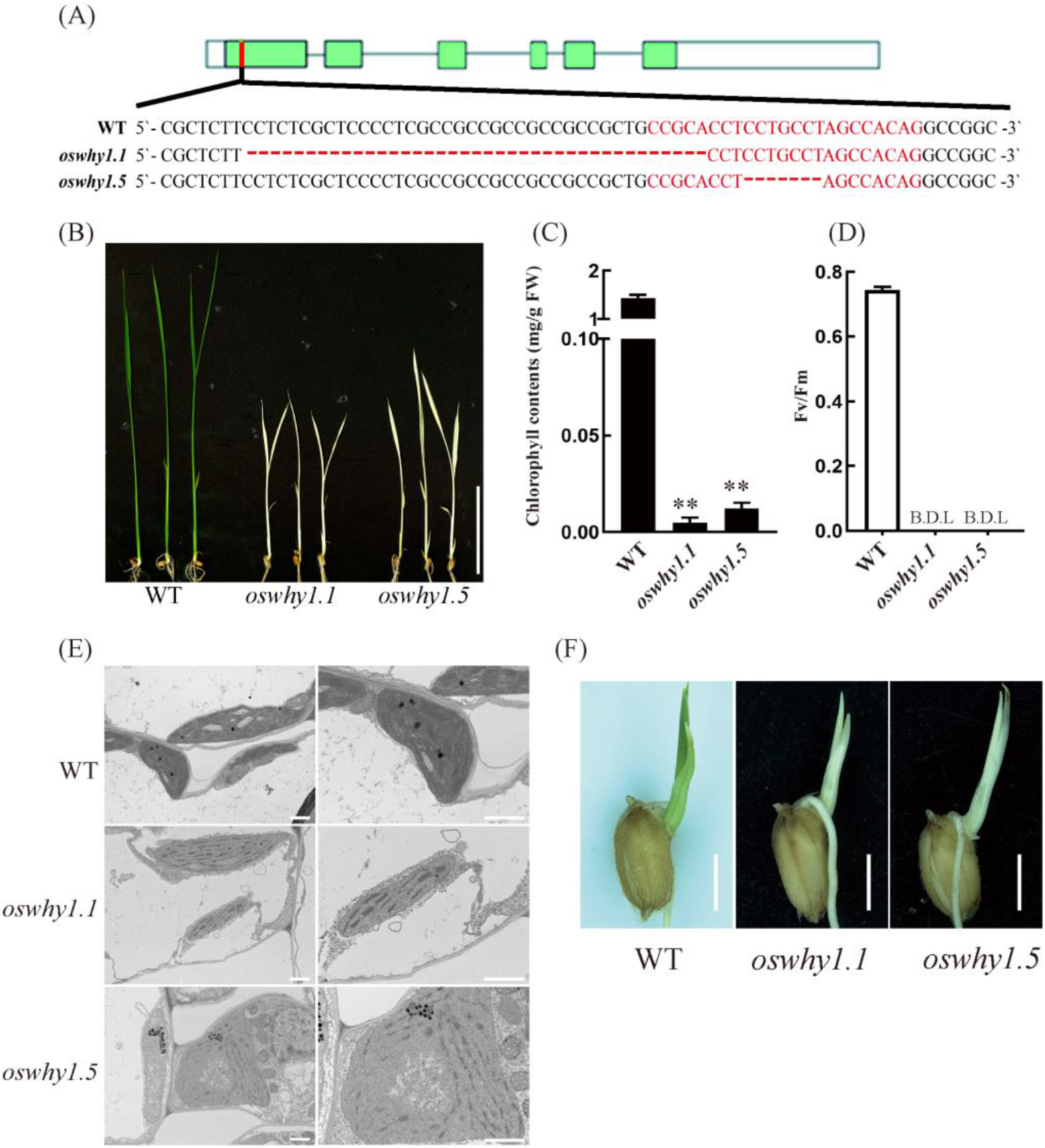
*oswhy1* mutants display albino seedling phenotype. (A) Structure of *OsWHY1* gene and the mutation sites in the *oswhy1* knockout lines. Green boxes indicate exons and the lines between them indicate introns. White boxes represent the 5′ and 3′ UTR. The 23-nt target site of the Cas9/sgRNA complex; the deletion sequences in *oswhy1.1* and *oswhy1.5* are highlighted in red. (B) Phenotypic comparison of 10-day-old seedlings of wild type (WT), *oswhy1* mutant plants under normal growth condition (Bar = 5.0 cm). (C, D) Chlorophyll contents (C) and relative photochemical efficiency of photosystem II (Fv/Fm) (D) of WT and the *oswhy1* mutants were measured using the second leaves of 10-day-old seedlings. Error bars indicate the mean ±SD (n=5). Significant differences of the chlorophyll levels normalized to WT were calculated using Student’s t-test (**, p < 0.01). B.D.L., below detection limit. FW, fresh weight. (E) Transmission electron microscope images of chloroplasts from WT and the *oswhy1* albinic mutant leaves. Bar = 2.0 μm. (F) Phenotype of WT and the two *oswhy1* mutants at third day after germination. Bar = 1.0 cm.

To further examine the albino leaf phenotype of *oswhy1*, ultrastructural features of chloroplasts from *oswhy1* mutant and WT plants were observed by transmission electron microscopy (TEM). The chloroplasts from the *oswhy1* seedlings presented disorganized lamellar structures with much smaller grana thylakoids and fewer grana lamella stacks (**Figure 1E**) compared to chloroplasts of WT, indicating that chloroplast development is severely impaired in the *oswhy1* leaves. These observations suggest that albino lethal phenotypes with disrupted chlorophyll accumulation and photosynthetic efficiency in *oswhy1* lines were probably due to aberrant chloroplast biogenesis.

Furthermore, seeds of *oswhy1* mutants appeared completely albino buds from the third day of germination (**Figure 1F**). Thus, OsWHY1 is required for early chloroplast development in rice.

In contrast, the generated overexpression lines of OE-OsWHY1-GFP exhibited stay-green leaves and maintained normal growth and development similar to the OE-GFP transgenic control plants under natural conditions (**Supplementary Figure S2**). These 3-week-old seedlings were subjected to darkness or intense light for 72 hours. It was observed that the photosynthetic efficiency of the leaves of the OE-OsWHY1-GFP overexpression lines was significantly higher than that of the OE-GFP control plants under high light conditions (**Supplementary Figure S2**), which implies that overexpression of OsWHY1 may enhance the chloroplast stability and thereby protect the photosynthetic system of the leaves from high light-induced damage.

### Altered transcript levels of plastid-encoded genes in *oswhy1* mutants

Since function of chloroplasts is regulated by both plastid and nuclear genomes, RT-qPCR was carried out to detect expression changes of several chloroplast-associated marker genes in the *oswhy1* mutants. The nuclear genes involved in chlorophyll synthesis, such as *CAO1* (encoding a chlorophyllide A oxygenase) and *HEMA1* (encoding a glutamyl-tRNA reductase), as well as photosynthesis-related genes, including *Cab1R* and *Cab2R* (encoding two light-harvesting chlorophyll a/b-binding proteins) were significantly down-regulated in the *oswhy1* lines compared with WT (**Figure 2A**). Transcriptome analysis also revealed that the majority of genes related to the photosynthesis pathway were strongly down-regulated in o*swhy1* albino seedlings compared to WT (**Supplementary Figure S4**). These results suggested that photosynthesis of *oswhy1* mutants is attenuated.

**Figure 2.**
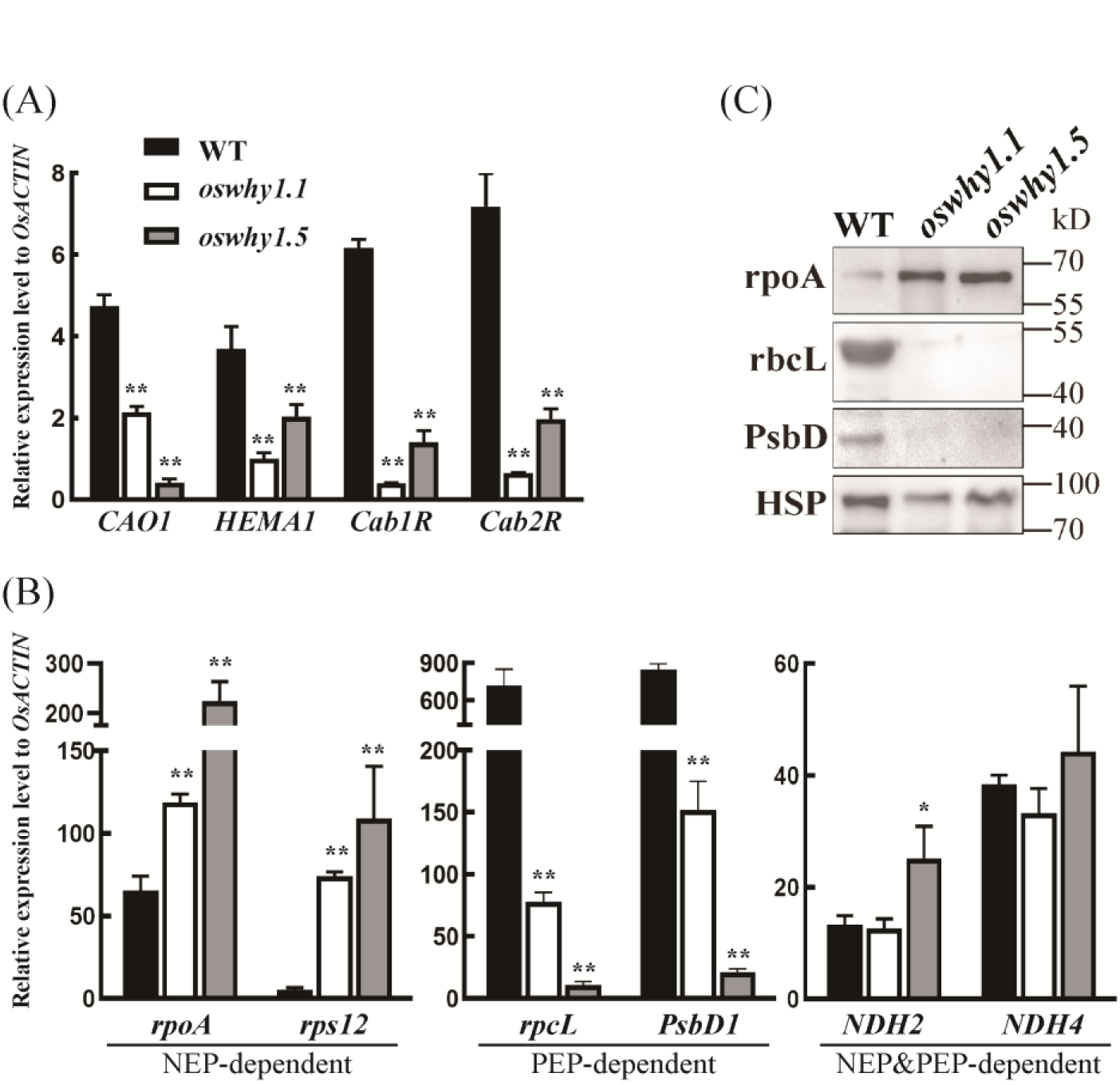
Expression changes of chloroplast-associated genes in the two *oswhy1* knockout lines. (A) Expression levels of genes related to chloroplast development and photosynthesis in WT and *oswhy1* mutant plants at the two-leaf stage. (B) Expression levels of NEP-dependent genes (*rpoA, rps12*), PEP-dependent genes (*rpcL, PsbD1*) as well as both NEP and PEP-dependent genes (*NDH2, NDH4*) in WT and *oswhy1* mutant plants at the two-leaf stage. Transcript levels are normalized relative to rice *OsACTIN* in each sample. Values represent the mean ± SD obtained from three biological replicates and three technical replicates. Significant differences of the expression level compared with WT were determined by Student’s t-test (*p < 0.05; **p < 0.01). (C) Western-blot analysis of chloroplast proteins in WT and *oswhy1* albino mutants. Proteins were extracted from 10-day-old seedlings, followed by probing immunoblots with antibodies against rpoA, rbcL and PsbD. HSP82 was used as an internal control.

Chloroplast biogenesis is closely related to transcription status of plastid genes, which is dependent on two types of plastid RNA polymerases: plastid-encoded plastid RNA polymerase (PEP) and nuclear-encoded plastid polymerase (NEP) (Borner et al., 2015). Analysis of RT-qPCR revealed that mRNA levels of PEP-dependent photosynthetic genes *PsbD1* and *rbcL* were strikingly decreased in the *oswhy1* mutants, while RNA polymerase gene *rpoA* and ribosomal protein gene *rps12* transcribed by NEP were strongly up-regulated in *oswhy1* knockout lines when compared to that in WT. Meanwhile, expression levels of NADH dehydrogenase genes *NDH2* and *NDH4* mutually transcribed by both PEP and NEP were slightly increased in the *oswhy1* mutants (**Figure 2B**). Western blot analysis also showed that the level of plastid protein ropA was increased in the *oswhy1* mutants relative to WT plants, whereas the plastid-encoded protein rbcL and PsbD were hard to be detected in the *oswhy1* knockout lines (**Figure 2C**), which is consistent with the transcript level analysis of *rpoA*, *rbcL* and *PsbD1*. Therefore, we propose that OsWHY1 is important for PEP complex activity and optimal expression of plastid genes during chloroplast development.

### OsWHY1 interacts with OsTRXz and OsMORFs in chloroplasts

In order to insight the molecular function of OsWHY1 in rice chloroplasts and whether OsWHY1 is involved in PEP complex activity, we immunoprecipitated the interacting candidates of OsWHY1 by GFP-nanotrap followed by MS assay (Supplementary Table S3). Interestingly, several PPR proteins potentially involved in chloroplast development and a plastid thioredoxin, OsTRXz, identified as a PEP-associated protein (Lv et al., 2017), were appeared in the list. Thus, some of them were selected to screen their interaction with OsWHY1 by yeast two hybrid (Y2H) and BiFC assays. The results revealed the association of OsWHY1 with OsTRXz, but no interaction between OsWHY1 and the obtained PPR proteins including OsPGL1 (**Figure 3A, Supplementary Figure S4**). In analogy to the subcellular distribution of OsWHY1-GFP, OsTRXz-GFP was enriched in rice chloroplasts with punctate localization pattern (**Supplementary Figure S4C**). Co-immunoprecipitation (Co-IP) experiment also confirmed the specific association of OsTRXz with OsWHY1, rather not with OsWHY2 (**Figure 3B**). Accordingly, these data indicated that OsWHY1 complexes with OsTRXz in chloroplasts, which possibly participates in the composition of a PEP complex.

**Figure 3.**
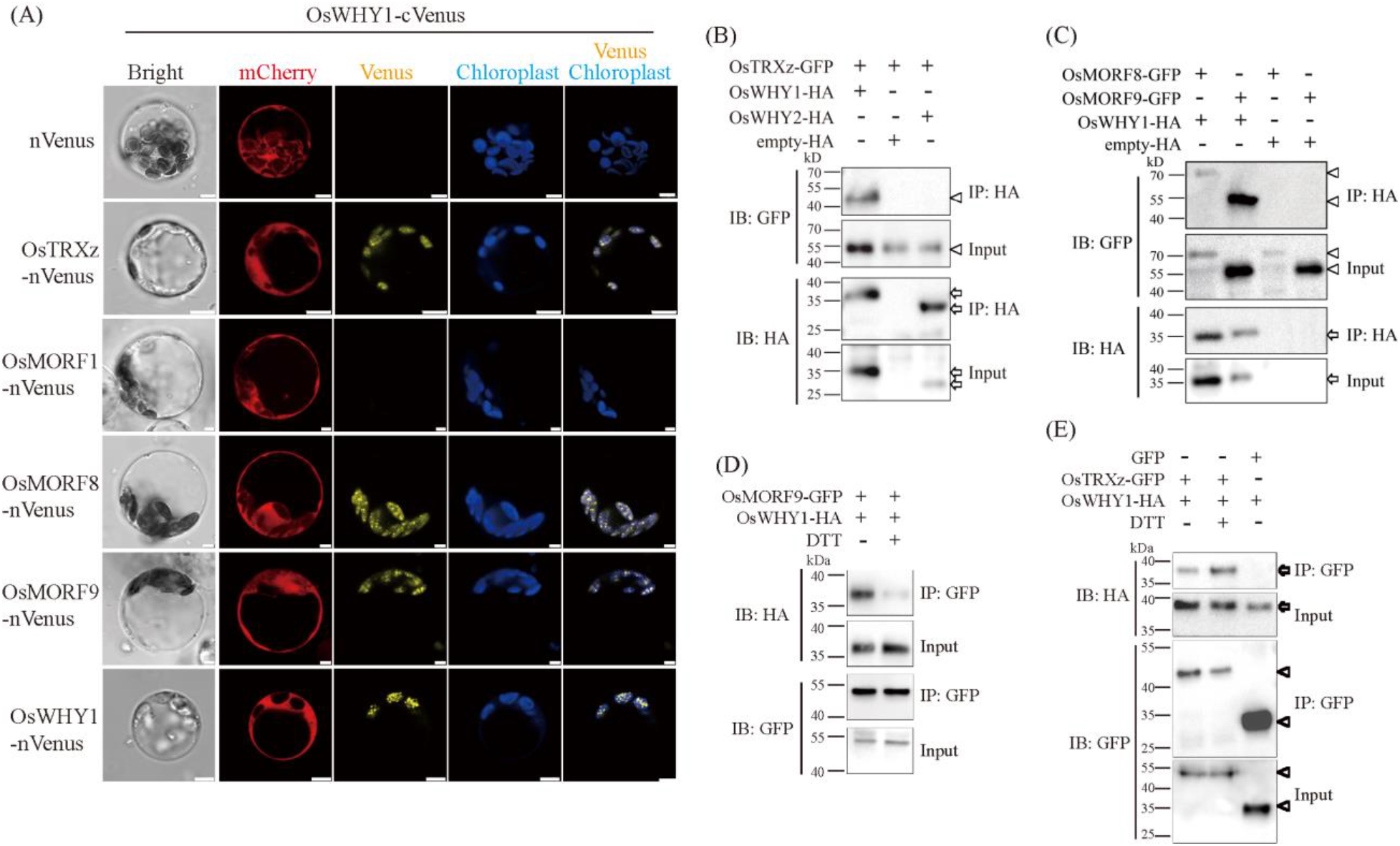
OsWHY1 interacts with OsTRXz as well as OsMORF proteins in rice chloroplasts. (A) The interaction detection of OsWHY1-cVenus with OsTRXz-nVenus, OsMORF1-nVenus, OsMORF8-nVenus, OsMORF9-nVenus and OsWHY1-nVenus by BiFC in rice protoplasts. Empty vector expressing nVenus is used as negative control. The independently expressed mCherry from the BiFC vector pRTVcVC is used as controls to monitor the construct transfection efficiency and gene expression in the transformed rice protoplasts. Bar = 3 µm. (B) *In vivo* Co- immunoprecipitation (Co-IP) assays showing that OsTRXz-GFP interacts with OsWHY1-HA, but not with OsWHY2-HA. Extracted proteins (input) from rice protoplasts co-expressing OsTRXz- GFP and OsWHY1-HA or OsWHY2-HA were subjected to immunoprecipitation (IP) with anti- HA affinity matrix followed by immunoblotting (IB) with anti-HA and anti-GFP antibodies. Arrows indicate HA-fusion proteins. Arrowheads indicate GFP-fusion proteins. The empty vector with HA tag (empty-HA) was used as a control. (C) *In vivo* Co-IP assays showing the interaction between OsWHY1-HA and OsMORF8-GFP or OsMORF9-GFP. Extracted proteins (input) from rice protoplasts co-expressing OsWHY1-HA and OsMORF8-GFP or OsMORF9-GFP were subjected to IP with anti-HA affinity matrix followed by immunoblotting (IB) with anti-HA and anti-GFP antibodies. Arrows indicate HA-fusion proteins. Arrowheads indicate GFP-fusion proteins. The empty vector with HA tag (empty-HA) was used as a control. (D) *In vivo* Co-IP assays showing that DTT treatment disturbs interaction between OsWHY1 and OsMORF9. Rice protoplasts co-expressing OsWHY1-HA and OsMORF9-GFP were treated with or without DTT, followed by protein extraction and IP with GFP-trap beads. (E) *In vivo* Co-IP assays showing that DTT treatment does not affect interaction between OsWHY1 and OsTRXz. Rice protoplasts co- expressing OsWHY1-HA and OsTRXz-GFP were treated with or without DTT, followed by protein extraction and IP with GFP-trap beads. Arrows indicate HA-fusion proteins. Arrowheads indicate GFP-fusion proteins.

It was recently reported that OsTRXz interacts with several plastid multiple organellar RNA editing factors (MORFs) and regulates RNA editing of many plastid-encoded genes (Wang et al., 2021b), which promotes the speculation about the relationship between OsWHY1 and OsMORFs. To this end, Y2H was first recruited to screen the interaction between OsWHY1 and obtained OsMORF members. It revealed that OsWHY1 can interact with OsMORF1 (LOC_Os11g11020) and OsMORF8 (LOC_Os09g33480), but not interact with OsMORF2 (LOC_Os06g02600), OsMORF3 (LOC_Os03g38490), OsMORF9 (LOC_Os08g04450) as well as OsWSP1 (LOC_Os04g51280) in yeast cells (**Supplementary Figure S5A**). However, it has been described that OsMORF8 and OsMORF9 can interact with each other to form heterodimer in the *in vitro* pull-down assays (Wang et al., 2021b). Considering that there are no plastids in yeast and limitations of heterologous protein-protein interaction in the Y2H system (Ferro and Trabalzini, 2013), we checked if OsMORF1, OsMORF8 and OsMORF9 were targeted in the same subcellular compartment as OsWHY1. Co-localization analysis performed in rice protoplasts revealed that OsMORF8-RFP and OsMORF9-RFP were distributed together with OsWHY1-GFP in chloroplasts, while OsMORF1-RFP mainly appeared in the cytoplasm and the nucleus and thus not co-localized with OsWHY1-GFP in chloroplasts (**Supplementary Figure S5B**). It implied that OsMORF1 might not be able to complex to OsWHY1 in rice cells. To test their interactions *in vivo*, BiFC assays were further carried out. Yellow fluorescence overlapped with autofluorescence of chlorophyll was observed when the combinations of OsWHY1-cVenus and OsMORF8-nVenus or OsMORF9-nVenus were co-expressed in rice protoplasts, while co-expression of OsWHY1-cVenus and OsMORF1-nVenus did not produce visible yellow fluorescent signals (**Figure 3A**), suggesting that OsWHY1 complexes with OsMORF8 and OsMORF9 in rice chloroplasts, but not with OsMORF1. Analogously, yellow fluorescent signals were also clearly detected in chloroplasts when OsWHY1-cVenus and OsWHY1-nVenus were co-transfected in rice protoplasts, indicating the homodimerization of OsWHY1 proteins, which is consistent with the reported tetramer structure of StWHY1 from potato (Desveaux et al., 2002; Cappadocia et al., 2012). Co-IP assays based on rice protoplast expression system further confirmed the association between OsWHY1 and OsMORF8 or OsMORF9 *in vivo* (**Figure 3C**).

As thioredoxins usually function as disulphide oxidoreductases to modulate the redox state and activities of their target proteins, we tested whether the OsWHY1-OsTRXz and OsWHY1- OsMORFs interactions are affected by redox. Co-IP assays showed that the association of OsWHY1 with OsMORF9 was significantly weakened in the co-transformed rice protoplasts under the treatment with DL- dithiothreitol (DTT), which is a strong reducing agent, compared to that without DTT treatment (**Figure 3D**). Intriguingly, *in vivo* interaction between OsWHY1 and OsTRXz seemed to be not influenced by DTT treatment (**Figure 3E**), implying that OsWHY1 may be physically targeted by OsTRXz in a redox-independent manner. These results indicated that the redox state is essential for the heterodimer formation of OsWHY1 and OsMORFs, which is similar to the interactions between the OsTRXz-targeted OsMORF proteins shown in the previous study (Wang et al., 2021b).

### OsWHY1 is required for effective *rps14-80* editing and *rpl2* splicing

Given the associations between OsWHY1 and OsTRXz as well as OsMORFs in chloroplasts, OsWHY1 might be involved in RNA modification of plastid transcripts. RT-PCR products followed by cDNA-bulk-sequencing analysis was employed to check whether mutation of *OsWHY1* affected editing efficiencies of 21 known RNA editing sites in chloroplast RNA (Corneille et al., 2000). It was found that the C-to-U editing efficiency of *rps14*-80 was dramatically reduced in both *oswhy1* knockout lines compared to that in WT (**Figure 4A, C**). The defect of editing at the *rps14*-80 site disturbed the conversion of the codon from UCA to UUA, which leaded to an amino acid replacement of leucine to serine. Other tested chloroplast genes were edited normally in the *oswhy1* mutants (**Supplementary Figure S6**). It indicated that OsWHY1 has impact on editing of *rps14*-C80 site, which would probably affect protein structure or function of RPS14.

**Figure 4.**
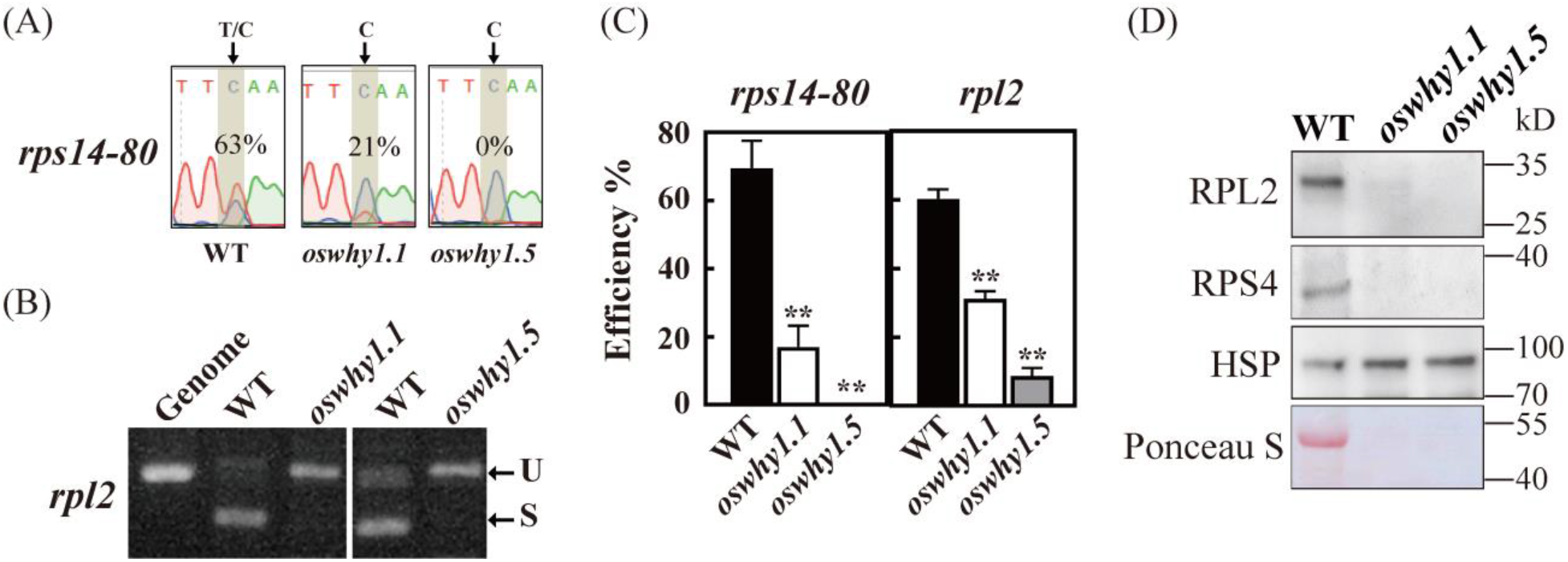
RNA editing and splicing analysis of the chloroplast transcripts in wild type (WT) and *oswhy1* mutant plants. Full datasets are presented in Supplementary Figure S6 and S7. (A) The editing levels of the *rps14* transcript at site 80 from WT and *oswhy1* leaves. The editing site of *rps14-80* is indicated by black arrows and the corresponding peaks are shaded in gray. The editing efficiency (C-T) is presented in the boxes. (B) Splicing analysis of the *rpl2* transcript in WT and the albino seedlings of *oswhy1* mutants. RT-PCR was performed with the RNA from leaves of 10-day-old seedlings. The amplification product from the genomic DNA (Genome) is shown on the left. Spliced (S) and unspliced (U) transcripts are shown at the right. (C) Quantitative analysis of the editing efficiency of *rps14-80* and the splicing efficiency *rpl2* in WT and the *oswhy1* mutants. The RNA editing levels of each site were determined by measuring the relative peak height of the nucleotide in sequence chromatograms and calculating the percentage of the height of “T” with respect to the sum of the height of “T” and “C”. The splicing efficiency of the transcript was obtained by measuring the band intensity of transcripts and calculating the percentage of band intensity of spliced transcript with respect to the sum of the band intensity of spliced and unspliced transcripts. Data are mean ±SD of four biological replicates. Significant differences of the editing or splicing efficiency in *oswhy1* mutants compared with WT were determined by Student’s t-test (**, p < 0.01). (D) Western-blot analysis of rice RPL2 and RPS4 protein in WT and *oswhy1* albino mutants. Total leaf proteins were extracted from 10-day-old seedlings, followed by probing immunoblots with RPL2 or RPS4-specific antiserum. HSP82 was used as an internal control. the Rubisco band was indicated by Ponceau S staining.

Many RNA binding proteins such as PPR and MORF proteins, are involved in both RNA editing and RNA splicing ((Sun et al., 2016; Zhao et al., 2020; Meng et al., 2024), we simultaneously detected plastid RNA splicing in the *oswhy1* mutants, RT-PCR was directly performed using specific primers flanking the introns and the lengths of PCR products were compared between WT and *oswhy1* mutants. One chloroplast transcript, *rpl2*, was spliced with much lower efficiency in both *oswhy1* homozygous lines compared to WT **(Figure 5B, C**). Splicing of other tested chloroplast genes was not evidently affected in the *oswhy1* mutants (**Supplementary Figure S6**). It suggested that OsWHY1 is important for RNA splicing of *rpl2* transcript, and probably influence protein translation and function of RPL2.

**Figure 5.**
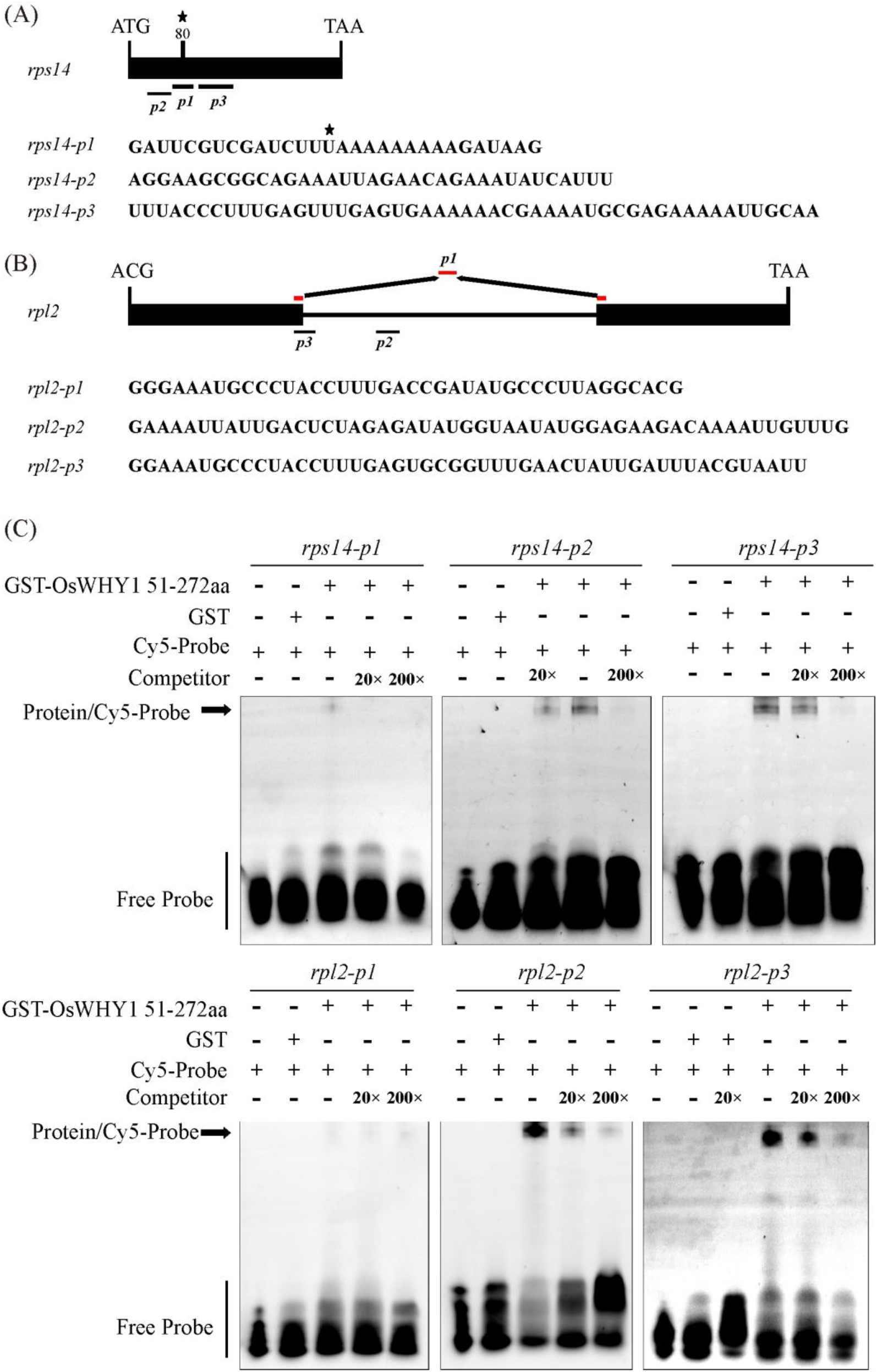
REMSA analysis of OsWHY1 protein with the *rps14* and *rpl2* oligoribonucleotides. (A) Sketch diagram and schematic sequences of the RNA probes *rps14-p1, rps14-p2* and *rps14- p3* on the corresponding pre-mRNA. The edited site *rps14-80* is indicated by asterisks. (B) Sketch diagram and schematic sequences of the RNA probes *rpl2-p1, rpl2-p2* and *rpl2-p3* on the corresponding pre-mRNA. The *rpl2-p1* probe (red) consists of exon sequences flanking the intron of the *rpl2* RNA precursor. (C) REMSAs showing that GST-OsWHY1 51-272aa binds to specific *rps14-p2, rps14-p3, rpl2-p2* and *rpl2-p3* RNA probes, but not to *rps14-p1* and *rpl2-p1* RNA probes. Cy5-probe, the RNA probe labeled with Cy5 at the 5’ end. Competitor, unlabeled RNA probe used as a competitor at a range of 20-fold or 200-fold excess concentrations for competitive REMSAs. GST protein was used as a negative control. GST-OsWHY1 51-272aa, GST, Cy5-labeled probes and competitors were present (+) or absent (-) in each reaction.

To further test if the stability of plastid ribosomal proteins was altered in the *oswhy1* mutants, we performed immunoblot analysis and showed that the level of ribosomal protein RPL2 and RPS4 were dramatically decreased in the *oswhy1* albino mutants relative to WT plant, and the RuBisCo band indicated by Ponceau staining was hardly detectable in both *oswhy1* lines, while the HSP expression levels in the WT and mutants did not significantly differ form one another **(Figure 4D)**.Thus, OsWHY1 is required for effective RNA processing of plastid *rps14* and *rpl2* transcripts as well as proper translation and accumulation of plastid-encoded ribosomal proteins.

### OsWHY1 binds to *rps14* and *rpl2* pre-mRNA

Due to the defective *rps14* editing and *rpl2* splicing in the *oswhy1* mutants, WHIRLY1 protein in Arabidopsis was existed in RNA-binding protein assembly (Lehniger et al., 2017), we anticipated that OsWHY1 would bind to certain sites in the pre-mRNAs of *rps14* and *rpl2*. The RNA electrophoresis mobility shift assay (REMSA) was designed to detect whether OsWHY1 is able to bind to RNA fragments of *rps14* and *rpl2*. The OsWHY1 truncated protein 51-272aa, minus the predicted plastid-targeted peptide, was fused with a GST tag for expression in *Escherichia coli*. (**Supplementary Figure S8A, B**). The purified recombinant GST-OsWHY1 51-272aa proteins were incubated with Cy5-labled RNA probes for *rps14.* These probes included 30 nucleotides (*rps14-p1*) surrounding the editing site of the *rps14*-80, as well as 35 (*rps14-p2*) and 50 (*rps14- p3*) nucleotides flanking the left and right sides of this fragment (**Figure 5A**). REMSA analysis showed that GST-OsWHY1 51-272aa bound strongly to the *rps14-p2* and *rps14-p3* RNA fragments, but weakly to *rps14-p1* (**Figure 5C, Supplementary Figure S8C**). No delayed band was observed when the RNA probe was incubated with the GST tag alone. Additionally, the specificity of this binding was confirmed using the same unlabeled oligoribonucleotide as a competitor **(Figure 5C, Supplementary Figure S8C**). Three RNA probes were designed and synthesized for the *rpl2* gene. These probes included 40 nucleotides (*rpl2-p1*) surrounding the splicing site of the mature *rpl2* RNA, 52 nucleotides (*rpl2-p2*) within the intronic region distal to the splicing site, and 49 nucleotides (*rpl2-p3*) surrounding the splicing site of the *rpl2* RNA precursor **(Figure 5B**). REMSA results revealed that GST-OsWHY1 51-272aa was able to bind the *rpl2-p2* and *rpl2-p3* RNA fragments, but not to *rpl2-p1*. Moreover, OsWHY1’s binding affinity to these pre-mRNAs decreased with increasing concentrations of competitive probes (Figure 5C, Supplementary Figure S8D). These findings demonstrated that OsWHY1 has RNA-binding activity and directly interacts with pre-mRNAs outside edited site of *rps14* and *rpl2*.

To clarify the molecular determinants of the association between OsWHY1 and the target RNA, a series of N-terminal or C-terminal deletion constructs of OsWHY1 were generated and fused to the GST tag for protein expression (**Figure 6A, B**). The purified proteins were incubated with RNA probes *rps14-p2, rps14-p3, rpl2-p2* or *rpl2-p3*. The REMSA results showed that with the exception of GST-OsWHY1 96-272aa, which like GST-OsWHY1 51-272aa can bind to these RNA fragments, the other three OsWHY1 mutants (GST-OsWHY1 128-272aa, GST-OsWHY1 51-235aa and GST-OsWHY1 97-235aa) lost their RNA binding activities (**Figure 6C, Supplementary Figure S9**). This suggests that the pNLS containing the conserved single-stranded nucleotide binding motif at the N-terminus of the Whirly domain as well as the potential autoregulatory domain (PAD) at the C-terminus of OsWHY1 play a key role in its RNA-binding ability, while the plastid transit peptide (PTP) and the putative transactivation domain (PTD) are not required for its RNA-binding activity.

**Figure 6.**
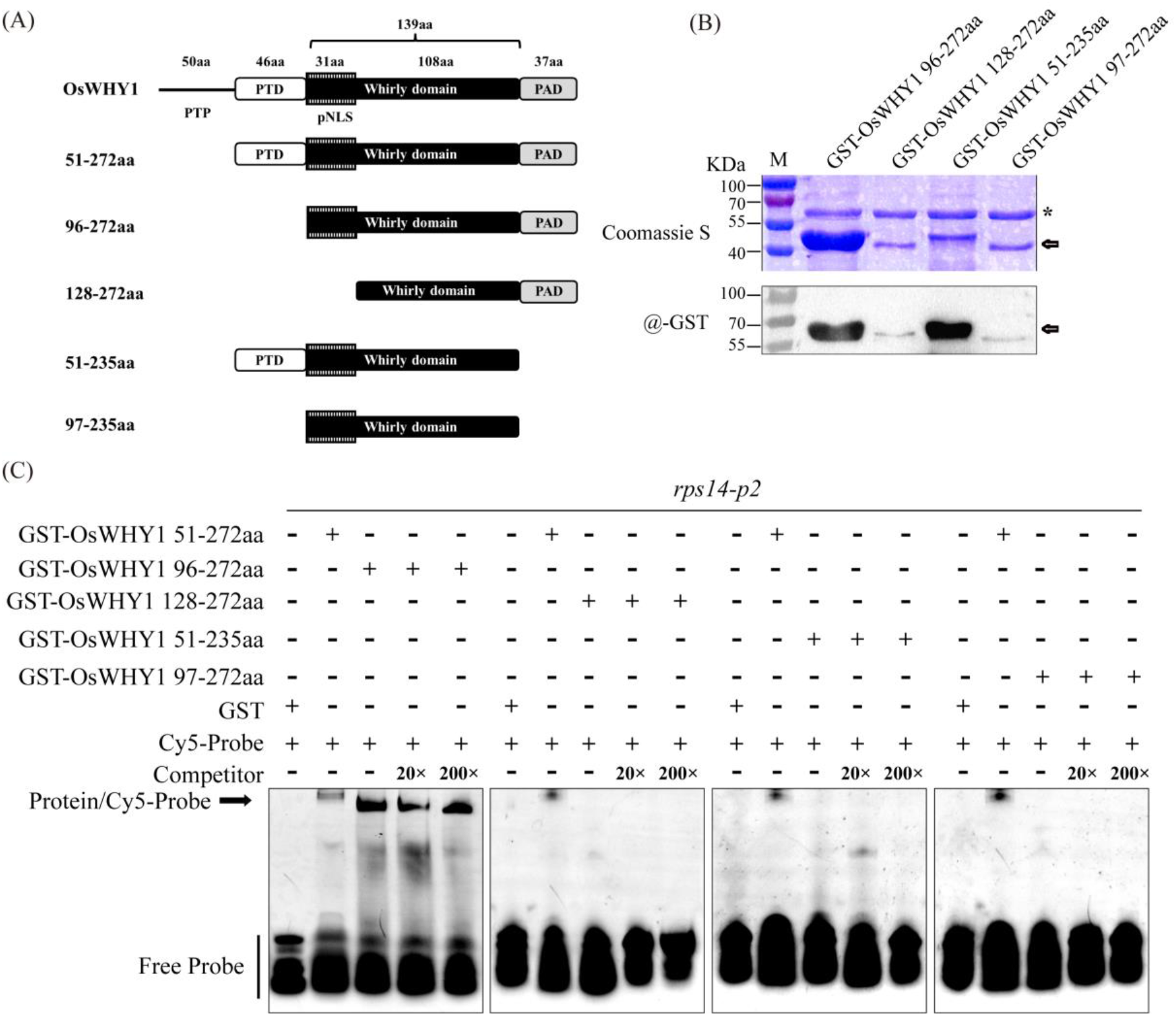
RNA binding activity analysis of OsWHY1 deletion mutants. (A) Schematic diagrams of the different *OsWHY1* deletion mutants used in this study. Numbers (left column) indicate positions of amino acid (aa) residues based on the full-length OsWHY1 protein sequence displaying the amino acid (aa) length of the predicted plastid transit peptide (PTP), putative transactivation domain (PTD), predicted nuclear localization signal (pNLS) containing the conserved KGKAAL motif for ssDNA-binding as well as putative autoregulatory domain (PAD). (B) The purified recombinant GST-OsWHY1 deletion mutants were isolated by SDS-PAGE, stained with Coomassie Brilliant Blue, and identified by Western blot. The corresponding GST-tagged OsWHY1 protein variants with expected molecular sizes are pointed out with an arrow. Non-specific band is pointed out with an asterisk. (C) REMSA analysis of OsWHY1 deletion mutants with the *rps14-p2* probe. Cy5-probe, the RNA probe labeled with Cy5 at the 5’end. Competitor, unlabeled RNA probe used as a competitor. GST protein was used as a negative control. GST-fused OsWHY1 deletion mutants, GST, Cy5-labeled probes and competitors were present (+) or absent (-) in each reaction.

### ROS accumulation is elevated in *oswhy1* mutants

The complex TRXz system in chloroplasts plays a central role in maintaining cellular redox balance. Due to the association of OsWHY1 and OsTRXz, the albino lethality of both *oswhy1* and *ostrxz* seedlings (He et al., 2018b; Wang et al., 2021b), we sought to examine in planta redox state of the *oswhy1* knockout lines. Production of superoxide radicals (O_2_^-^) and H_2_O_2_ in leaves of two-week-old seedlings was measured using NBT and DAB staining methods, respectively. Both *oswhy1* mutants showed obviously higher O_2_^-^ and H_2_O_2_ levels compared with the WT, which was reflected by precipitation of more densely blue formazan or reddish-brown formazan in *oswhy1* albino leaves than in WT (**Figure 7A**). Consistent with the DAB staining, the content of H_2_O_2_ was significantly increased in both *oswhy1* lines than in WT plants (**Figure 7B**), indicating that loss of function of *OsWHY1* resulted in over-production of ROS in rice leaves.

**Figure 7.**
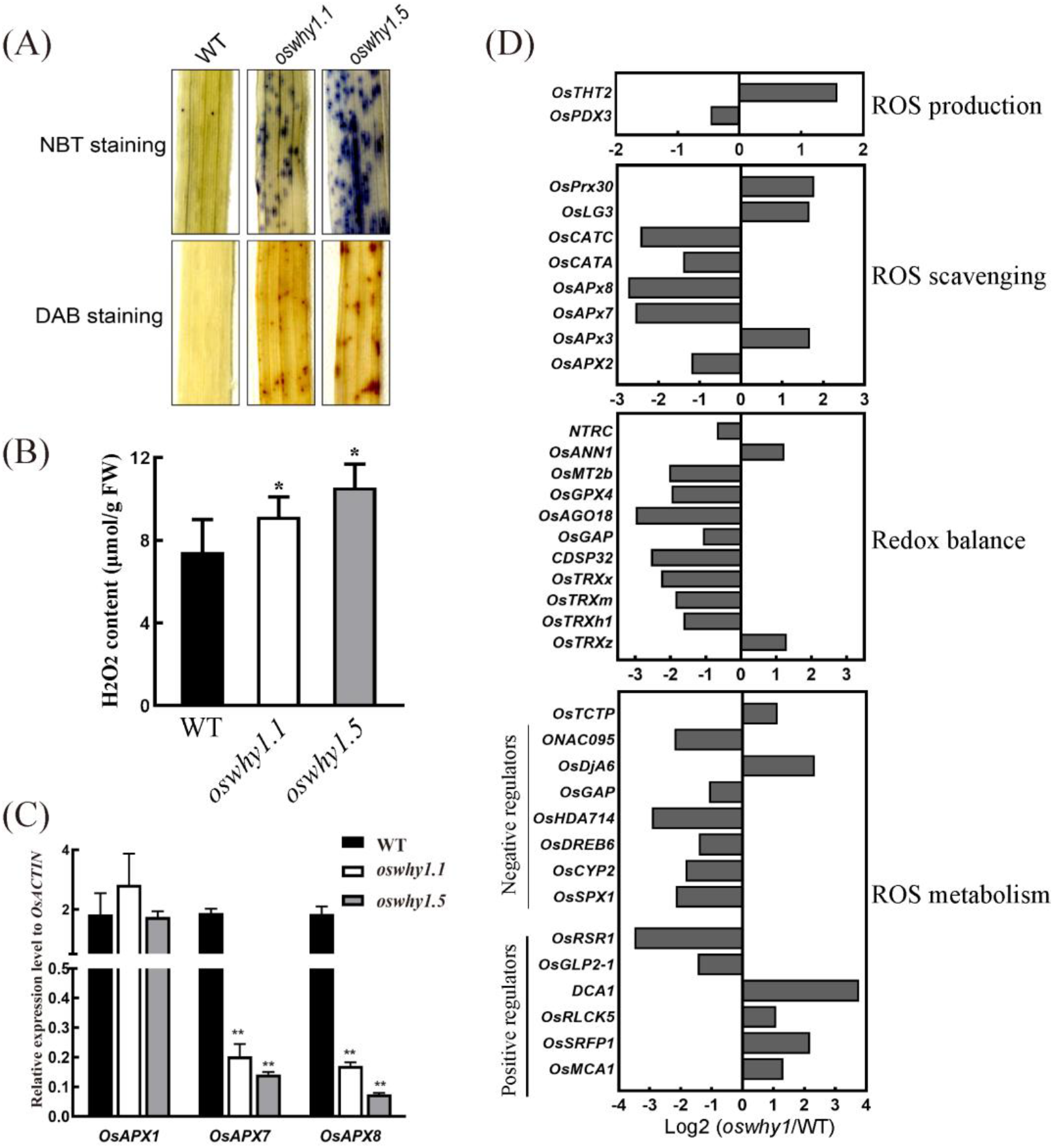
The *oswhy1* mutants showed elevated ROS accumulation. (A) NBT and DAB staining of the leaves from WT and *oswhy1* mutants. (B) Quantitative measurement of H_2_O_2_ from leaves of 10-day-old seedlings of WT and oswhy1 mutants (n=3 repeats.). (C) Relative expression levels of several ROS detoxification-related genes encoding ascorbate peroxidase (APXs) in WT and oswhy1 mutants. RNA was extracted from leaves of 10-day-old seedlings. Transcript levels are normalized relative to rice *OsACTIN* in each sample. Values represent the mean ±SD obtained from three biological replicates. Significant differences of the expression level compared with WT were determined by Student’s t-test (**, p < 0.01). (D) Transcripts of differentially expressed genes (DEGs) related to ROS metabolism and redox balance by RNA-seq analysis of *oswhy1* and WT. Log2 (*oswhy1*/WT) indicates the log2 ratio of mRNA levels in *oswhy1* mutant compared to WT.

To afford a molecular explanation for the effect of *OsWHY1* on H_2_O_2_ homeostasis, the transcript amounts of ROS associated genes between *oswhy1* and WT plants were compared by RT-qPCR and RNA-seq analysis. The expression of several genes associated with ROS degradation, including the genes encoding ascorbate peroxidase (APX) or catalase (CAT), were repressed in *oswhy1* (**Figure 7C, D**). Interestingly, the majority of the genes belonging to the GO termed cellular redox balance (GO:0045454) based on the China Rice Data Center (https://ricedata.cn/ontology/), such as *OsTRXx*, *OsTRXm*, *OsTRXh1* and *CDSP32* encoding a thioredoxin-like protein, were dramatically down-regulated in *oswhy1* (**Figure 7D**), pointing to the broken redox balance in *oswhy1* mutants. Thus, the mRNA levels of many stress responsive genes, such as *ONAC095, OsDREB6, OsCYP2* and *DCA1,* implicated in the ROS metabolic process were substantially changed in *oswhy1* compared with WT (**Figure 7D**). These results suggested that the albino phenotype of *oswhy1* mutants was caused not only by damaged development of chloroplasts but also by excessive ROS accumulation in leaves.

### Plastid signals mediating nuclear transcriptome reprograming are occured in *oswhy1* mutant

Due to the fact that most of the proteins required for chloroplast development and function are encoded by nuclear genes, changes in chloroplast functional status often give rise to signals regulating nuclear gene transcription *via* retrograde signaling. The phenomenon of retrograde signaling was first noticed in studies of albino barley (*Hordeum vulgare*) and maize (*Zea mays*) seedlings, which accumulated or reduced mRNA levels from several Photosynthesis-Associated Nuclear Genes (*PhANGs*) (Mayfield and Taylor, 1984; Batschauer et al., 1986; Burgess and Taylor, 1987; Rapp and Mullet, 1991; Hess et al., 1994; Borner, 2017; Kendrick et al., 2022). The important plastid signals are ROS dynamics and redox state (Chan et al., 2016; Kleine and Leister, 2016; Brunkard and Burch-Smith, 2018). Considering the interaction between OsWHY1 and OsTRXz as well as its effect on redox balance, we further sought to assess the function of OsWHY1 in nuclear gene expression reprograming.

To this end, we analyzed transcriptomes of non-photosynthetic albino rice *oswhy*1 mutant and compared them to transcriptomes of stages of normal leaf development. We identified ∼3,718 differentially expressed genes (DEGs) with 1871 DEGs upregulated, 1847 DEGs down-regulated (|log_2_FC|≥1, P value <0.01) (**Supplementary Table S1**), which fall into top 20 major categories based on the polarity and mutant-specificity of the change. Downregulated genes were enriched in metabolic and photosynthesis pathway for functions in photosynthesis **(Figure 8A)**, whereas upregulated genes were enriched in ribosome and aminoacyl-tRNA biosynthesis pathway for functions in chloroplast biogenesis and cytosolic translation (**Figure 8B**). GO analysis further showed that the 56 of 78 DEGs related to chloroplast development caterogy were upregulated in *oswhy1* mutant, including regulatory genes related to plastid ribosome function (*WP1, RLS, RPL21*), PEP activity (*pTAC3, OsTRXz, OsFLN1Os*), plastid RNA editing or splicing (*WSP1*, *PPR, MORF8* and *MORF9*) as well as plastid proteostasis (*OsClpP, OsBE1, OsChlD, YL1*) (**Figure 8C**); 21 of the 78 DEGs were downregulated in *oswhy1* mutant, including the genes involved in chloroplast structural development (*OsHCF222, OsWSL9, OsHAP3C, OsGIC*), chlorophyll biogenesis for photosynthesis (*OsABCI7, OsYGL8, OsGLK1*) as well as nutrient metabolism (*OsACS6, OsHAD1, OsAld-Y*) (**Figure 8D**); 33 of 35 DEGs related to *PhANGs* were significantly downregulated in *oswhy1* mutant (**Figure 8D**). RT-qPCR analysis also confirmed that transcript levels of *OsMORF8* and *OsMORF9* were strongly increased in both *oswhy1* knockout lines, while the transcription of three *PhANGs* (*OsPsbS, OsPsb28* and *ATP synthase B*) were remarkably reduced in *oswhy1* mutants (**Figure 8E**). This result reveals that loss-of-function of OsWHY1 promotes nuclear transcriptional reprogramming and multiple retrograde signals mediating plastid transcription, plastid RNA processing, as well as plastid protein synthesis and degradation, which is reflected by upregulation of regulatory genes associated to early plastid biogenesis and downregulation of chloroplast structural genes as well as functional PhANGs, leading to a defect in chloroplast development.

**Figure 8.**
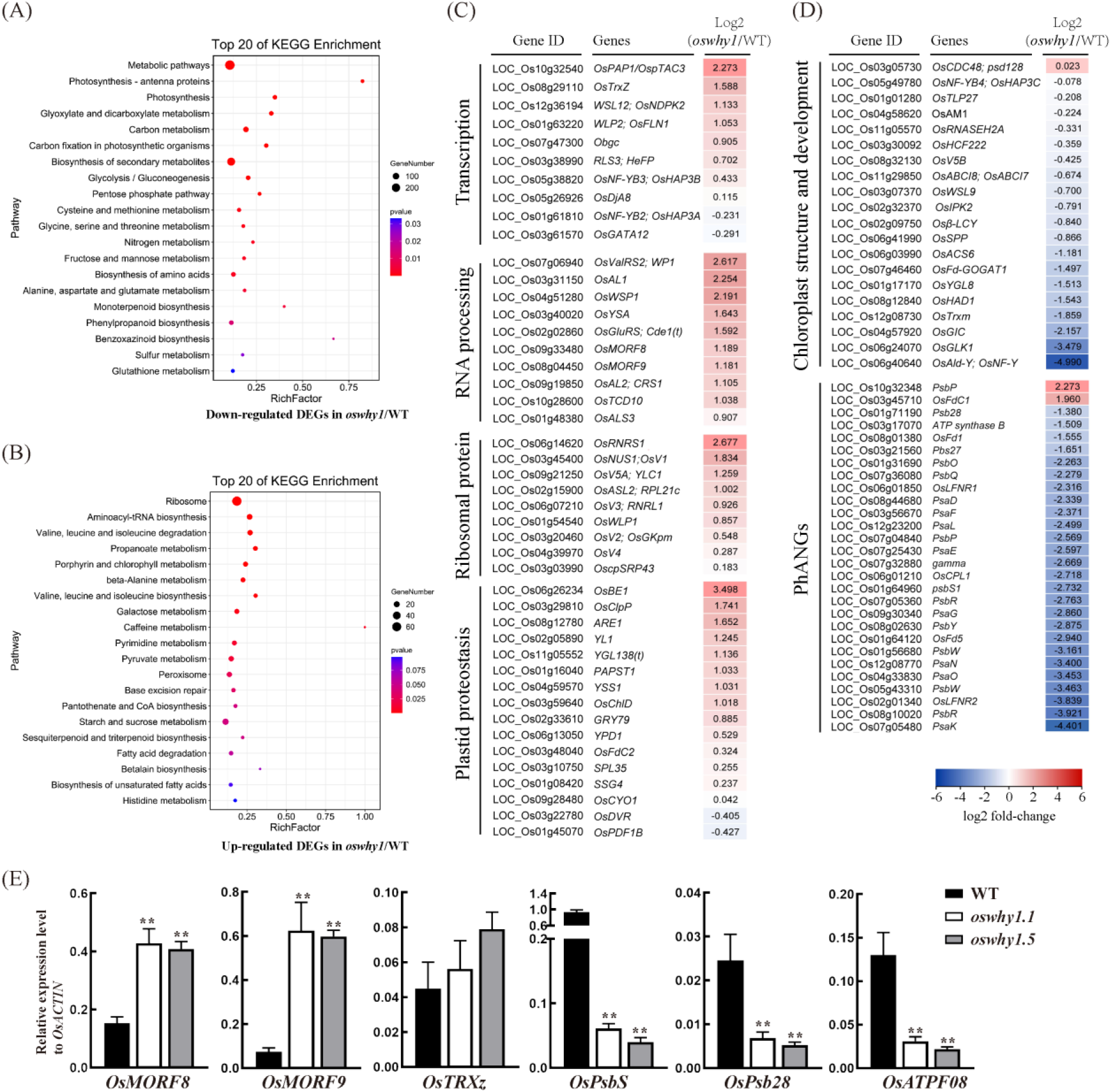
Transcriptome analysis of *oswhy1* and WT. (A) KEGG pathway enrichment analysis of differentially expressed genes (DEGs) lower expressed (Log2Fold change ≤ 1) in *oswhy1* mutant plants relative to WT plants. The top 20 items are displayed in the graph. (B) KEGG pathway enrichment analysis of DEGs higher expressed (Log2Fold change ≥ 1) in *oswhy1* relative to WT. The top 20 items are displayed in the graph. (C, D) Expression heatmaps of selected chloroplast development and photosynthesis-associated genes in *oswhy1* compared to WT. Log2 (*oswhy1*/WT) indicates the log2 ratio of mRNA levels in *oswhy1* mutant compared to WT. (C) Nuclear genes involved in chloroplast biogenesis: transcriptional regulation, RNA processing, plastid translation (ribosomal proteins) and plastid proteostasis. (D) Nuclear genes involved in chloroplast development and function: chloroplast structure and PhANGs. (E) Relative expression of several nuclear genes related to plastid biogenesis and PhANGs was examined in WT and *oswhy1* mutant plants at the two-leaf stage. Transcript levels are normalized relative to rice *OsACTIN* in each sample. Values represent the mean ± SD obtained from three biological replicates. Significant differences of the expression level compared with WT were determined by Student’s t-test (**p < 0.01).

### Involvement of OsMORF9 and OsTRXz in Chloroplast Development

Given that OsWHY1, OsMORF9 and OsTRXz can interact with each other in rice chloroplasts, it is speculated that these proteins form a functional complex in chloroplast development. To this end, we generated knockout mutants of *OsMORF9* in ZH11 by CRISPR/Cas9 genome editing. Two homozygous *osmorf9* mutants were identified, named *osmorf9.1* and *osmorf9.2*, with *osmorf9.1* harboring a 6-bp deletion from position 138 to 143 and *osmorf9.2* harboring a 7-bp substitution from position 137 to 478 in the target exon region (**Figure 9A**). Similar to the phenotype observed in the *oswhy1* mutants, both *osmor9* mutant lines showed an albino seedling phenotype (**Figure 9B**). We also analyzed the *white panicle2* mutant (*wp2*), which harbors a mutation in the *OsTRXz* gene (Wang et al., 2021b). The *wp2* seedlings exhibited an albino phenotype when subjected to high temperatures, maintaining this phenotype upon return to normal growth conditions (**Figure 9C**). This consistency with previous findings (Wang et al., 2021b) supports the role of OsTRXz in chloroplast function.

**Figure 9.**
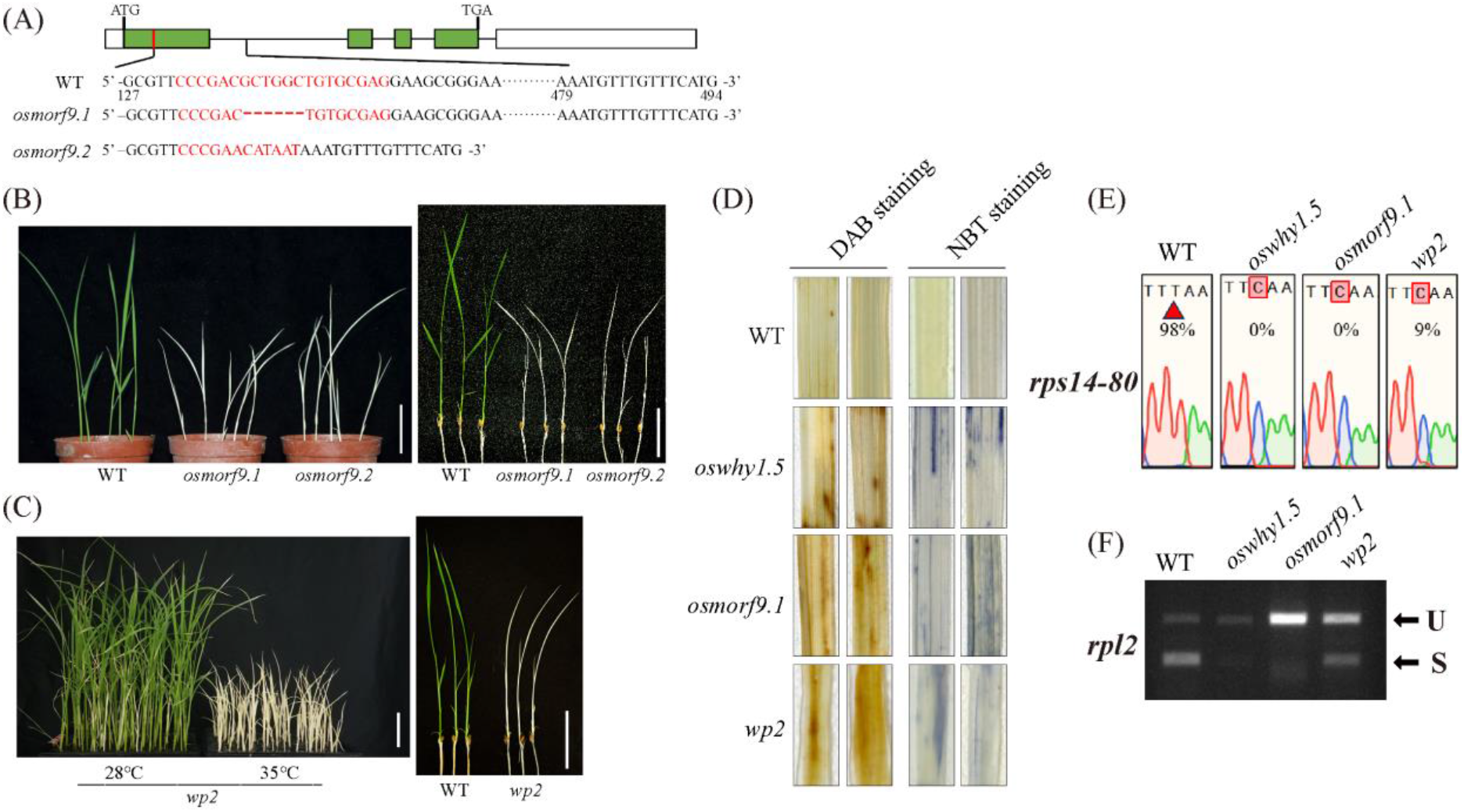
Phenotypic and molecular analysis of *osmorf9* and *wp2/ostrxz* mutants. (A) Schematic representation of the *OsMORF9* gene structure and mutation sites in *osmorf9* knockout lines. Green boxes denote exons, lines between them indicate introns, and white boxes represent the 5′ and 3′ UTRs. The 20-nt target site, a 6bp deletion in *osmorf9.1*, and a 7bp substitution from position 137 to 478 are highlighted in red. (B) Phenotype of *osmorf9* albino seedlings; bar = 3 cm. (C) Phenotype of *wp2* mutant (*ostrxz* mutant) seedlings under 28°C and 35°C growth conditions; bar = 3 cm. (D) NBT and DAB staining of the leaves from WT green seedlings and albino seedlings of *oswhy1*, *osmorf9*, or *wp2*. (E, F) RNA editing efficiency at the *rps14-80* site (E) and RNA splicing efficiency of *rpl2* (F) in *oswhy1*, *osmorf9* and *wp2* albino seedlings.

To further understand the impact of these mutations, NBT and DAB staining were conducted on the leaves of two-week-old *oswhy1.5, osmorf9.1, wp2* albino seedlings, alongside WT green seedlings. The results showed increased accumulation of O_2_^-^ and H_2_O_2_ in the leaves of the albino mutants compared to the WT (**Figure 9D**), indicating disrupted chloroplast development and redox homeostasis. Additionally, RNA editing and splicing assays demonstrated significantly reduced editing efficiency at the *rps14-80* site and impaired splicing efficiency of the *rpl2* transcript in the albino seedlings of *oswhy1, osmorf9* as well as *wp2* mutants (**Figure 9E, F**). These findings suggest that the OsWHY1-OsTRXz-OsMORF9 complex is crucial for the RNA modification of plastid genes, thereby influencing chloroplast development and maintaining cellular ROS balance.

## Discussion

Chloroplast biogenesis is coordinately regulated by plastidic and nuclear factors, since many nuclear genes encoding plastid protein plays a key role in chloroplast development. One member of the WHIRLY protein family, WHY1 has been reported to have dual locations in plastids and nuclei and to play diverse functions such as plant senescence and stress response by coupling with DNA or RNA in both compartments (Krause et al., 2005; Grabowski et al., 2008; Marechal et al., 2009; Ren et al., 2017; Duan et al., 2020; Wang et al., 2021a). However, in various species it exhibited different functions (Prikryl et al., 2008; Marechal et al., 2009; Krupinska et al., 2014; Ren et al., 2017). Our study shows that OsWHY1 is mainly located in chloroplasts and plays a critical role in early chloroplast development through complexation with OsTRXz and multiple OsMORF proteins. The findings highlight OsWHY1’s involvement in RNA processing and redox balance, providing new insights into the molecular mechanisms governing chloroplast biogenesis and stability (**Figure 10**).

**Figure 10.**
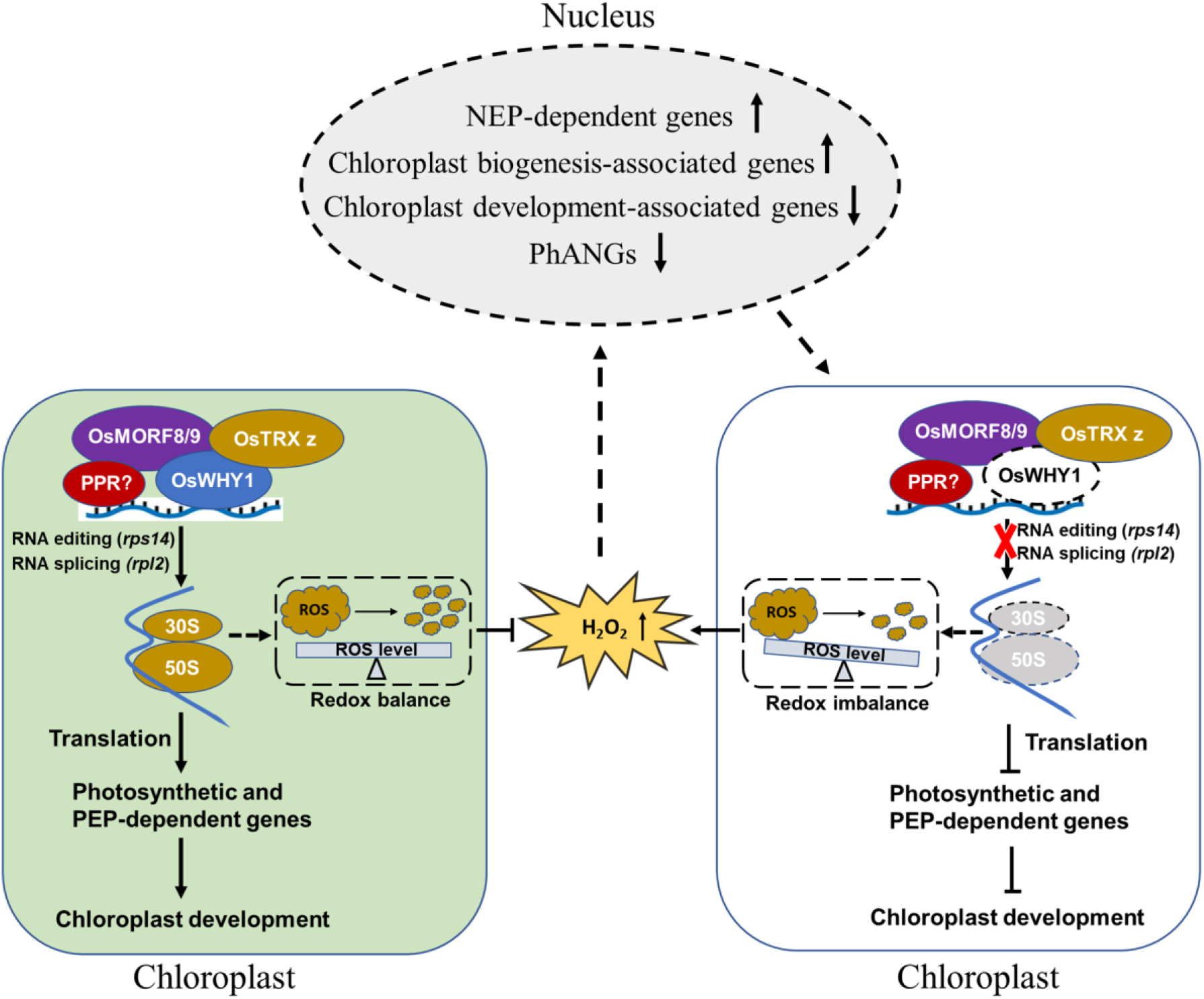
A schematic diagram illustrating the possible mechanism underlying the albinic phenotype of *oswhy1* mutants. The OsWHY1/OsTRXz/OsMORFs complex is involved in plastid RNA processing and is required for chloroplast development in rice. The main characteristic of the *oswhy1* mutant is the deficiency in RNA editing and splicing, and the remaining defects including decreased chlorophyll content, changed gene expression as well as redox imbalance are secondary. The plastid ribosome genes *rps14* and *rpl2* fail to be normally edited or spliced in the *oswhy1* mutants, which results in defective RPL2 expression and ribosome biogenesis, which in turn probably affects PEP component translation and redox balance. Thus, hampered transcription and translation of plastid genes lead to aberrant chloroplast development and defect in photosynthesis. On the other side, elevated ROS level can give rise to nuclear transcriptome reprogramming. The PPR? indicates that there may be an unidentified PPR family member directly involved into *rps14* editing or *rpl2* splicing.

### The chloroplast-localized OsWHY1 is essential for early stages of chloroplast biogenesis

Although WHY proteins are highly conserved across higher plants and well-characterized in the last two decades, the functions of WHYs seem not to be fully understood, as exemplified by the phenotypic inconsistency of *why1* mutants in different plants. Single knockout mutant of *Atwhy1* or *Atwhy3* showed an early senescence or no apparent phenotype under standard growth conditions, while a small ratio of the *Atwhy1why3* double mutants appeared with variegated green/white/yellow leaves, indicating the involvement of the two plastidic AtWHY proteins in chloroplast development (Marechal et al., 2009; Lepage et al., 2013). Knockdown lines of *WHY1* in barley, tomato and cassava are phenotypically similar to wild-type (WT) plants when grown under optimal conditions (Comadira et al., 2015; Zhuang et al., 2019; Yan et al., 2020). In contrast, loss of function mutant of *ZmWHY1* in maize results in severe albino phenotypes and seedling lethality due to deficiency in *atpF* intron splicing and plastid ribosomes (Prikryl et al., 2008). Herein, CRISPR/Cas9-edited *oswhy1* mutant lines exhibit the same phenotype as the homozygous *Zmwhy1-1* mutants as well as the recent published work on *OsWHY1* (**Figure 1**) (Prikryl et al., 2008; Qiu et al., 2022). Our findings, however, indicated that OsWHY1 only impacted the splicing of the plastid-encoded *rpl2* transcripts (**Figure 4, Supplemental Figure S7**), which was inconsistent with ZmWHY1 and HvWHY1 (Prikryl et al., 2008; Melonek et al., 2010). The splicing efficiency of the plastid gene *atpF* in the two independent *oswhy1* mutant lines that we found here did not differ substantially from the wild type. The *ndhA* gene’s splicing efficiency was normal in the *oswhy1.5*, despite being slightly lower in the *oswhy1.1* (**Supplemental Figure S7**), which is less consistent with the effect of *ndhA* gene splicing in the *OsWHY1* knockout mutants demonstrated in earlier studies (Qiu et al., 2022). In line with the previous findings, RNA editing efficiency at the *rps14-80* site was significantly reduced in all *oswhy1* mutants (**Figure 4, Supplemental Figure S6**). Thus, it reveals that the *oswhy1* albino seedling phenotype is not linked to the splicing of the *atpF* and *ndhA* genes, but rather most likely to post-transcriptional modifications of the two PEP-dependent plastid genes *rpl2* and *rps14*, which encode the 50S ribosomal protein RPL2 and the 30S ribosomal protein RPS14, respectively.

RPL2 and RPS14 are crucial components of the chloroplast translational apparatus, and are required for plastid translation in tobacco and ribosomal assembly in *E.coli* (Ahlert et al., 2003; Shoji et al., 2011; Tiller and Bock, 2014). Defective *rpl2* splicing accompanied by decreased accumulation of RPL2 has been found in different rice mutants displaying white-striped seedling phenotype and abnormal chloroplast development, like *wsl, wsl4* and *wsl9* (Tan et al., 2014; Wang et al., 2017; Zhu et al., 2020). Impaired editing events in *rps14* were observed in different Arabidopsis mutants showing albino phenotype or embryonic lethality, such as *ppi2-2*, *ecd1* and *emb2261* (Kakizaki et al., 2012; Jiang et al., 2018; Sun et al., 2018). The RNA processing of *rps14* and *rpl2* is essential for the synthesis of functional RPS14 and RPL2 proteins, which are crucial for ribosome biogenesis and plastid translation. In fact, we also observed that several plastid-encoded ribosomal proteins, such as RPL2 and RPS4, as well as functional proteins like rbcL and PsbD, are extremely low expressed in the *oswhy1* mutants (**Figure 2**; **Figure 4**). Hence, we argue that OsWHY1’s involvement in RNA processing of *rps14* and *rpl2*, which is crucial for normal protein synthesis in chloroplasts, is a likely explanation for the *oswhy1* phenotype. We also generated *RNAi OsWHY1* knockdown lines, most of which did not show obvious phenotypes, except some individuals displaying albino phenotype (**Supplementary Figure S10**). It seemed possible that decreased expression of OsWHY1 by posttranscriptional gene silencing method may not completely block the translation of OsWHY1, leading to comprised but still enough OsWHY1 working in chloroplasts. This finding implies that OsWHY1 might be required for early stages of chloroplast biogenesis in rice in a dose dependent manner.

### OsWHY1 interacting with OsTRXz-OsMORFs provides an RNA-binding link between editosomes and the target transcripts

In addition to the defective RNA processing, our results show that PEP activity is disrupted in the *oswhy1* mutants, as the transcript levels of the NEP-dependent genes were up-regulated and the transcripts of the PEP-dependent plastid genes were significantly low in both *oswhy1* mutant lines (**Figure 2**). In Arabidopsis, WHY1 was found to a part of the isolated plastid transcriptionally active chromosome (pTAC) complex (Pfalz et al., 2006), implying that WHY1 is a PEP-associated protein (PAP). It has been well documented that many PPR family members are RNA-binding proteins involved in RNA processing (Sun et al., 2016; Zhao et al., 2020; Meng et al., 2024). Thus, in the results of IP-MS assays of OsWHY1-GFP, we focused on the proteins that may be implicated in PEP complex and RNA metabolism. Although several selected PPR proteins did not interact with OsWHY1, a chloroplast thioredoxin-like protein (OsTRXz) was shown to interact with OsWHY1 in plastids (**Figure 3, Supplementary Figure S4**), which is consistent with the recent publication on OsWHY1 (Qiu et al., 2022). As a part of pTAC, TRXz affects PEP activity through interaction with different chloroplast proteins, including FLN1, FLN2, TSV, PRIN2, and CHLI (Pfalz et al., 2006; Arsova et al., 2010; Schroter et al., 2010; Zhang et al., 2015; Chang et al., 2017; Lv et al., 2017; Sun et al., 2017; Diaz et al., 2018; He et al., 2018b). The formation of OsWHY1-OsTRXz complex suggests one role of OsWHY1 in maintaining PEP activity for proper transcription of plastid genes.

On the other side, OsTRXz functions in RNA editing by interacting with all chloroplast OsMORFs, which are well-known as RNA editing factor interacting proteins (RIPs) due to their participation in PPR-RNA complexes (Xiao et al., 2018a; Cui et al., 2019; Wang et al., 2021b). To find out direct molecular evidence linking OsWHY1 to the RNA editing machinery, we screened and identified the association between OsWHY1 and OsMORF8 or OsMORF9 in rice chloroplasts (**Figure 3, Supplementary Figure S5**), which emphasizes the involvement of OsWHY1 in plastid RNA processing. OsTRXz, known for its redox-regulating functions by reducing the target proteins, modulates the redox state of OsMORFs, which is essential for the dynamic assembly of activated editosomes (Wang et al., 2021b). In our study, it is found that the interaction between OsWHY1 and OsMORF proteins is redox-dependent, whereas the interaction between OsWHY1 and OsTRXz is redox-independent (**Figure 3**). This suggests that OsWHY1, along with OsMORF8 and OsMORF9, are substrates for OsTRXz’s redox modulation. The redox state, regulated by OsTRXz, influences the interaction among these proteins, which is critical for the formation of functional RNA editing complexes. This regulatory mechanism underscores the importance of redox balance in chloroplast RNA processing and stability. Possibly due to this unstable interaction between OsWHY1 and OsMORFs, no OsMORF protein-related peptide signals were shown in the IP-MS results, whereas OsTRXz peptide signals were clear. The similar phenotypic abnormalities observed in the *oswhy1*, *osmorf9*, and *wp2/ostrxz* mutants further support the functional association between OsWHY1, OsMORF9, and OsTRXz. These mutants exhibit disrupted chloroplast development and defective RNA processing (**Figure 9**), confirming that OsWHY1/OsTRX z/OsMORF9 forms a regulatory complex essential for RNA modification of plastid genes and the maintenance of chloroplast integrity.

Potato StWHY1 was first identified as a transcription factor in the early 2000s when it was recognized for its DNA-binding activity (Despres et al., 1995; Desveaux et al., 2000), and subsequent studies further revealed that WHY1 in maize and barley exhibits significant RNA-binding activity and is associated with intron-containing plastid RNAs, including the *atpF* and *rpl16* transcripts (Prikryl et al., 2008; Melonek et al., 2010). However, the precise molecular mechanism underlying the interaction between WHY1 and targe RNAs remains obscure. In the present study, the RNA-binding activity of OsWHY1 was firstly confirmed by REMSA assays, and structure-function analysis revealed that OsWHY1 loses its RNA-binding activity when its N-terminal putative nucleus localization signal (pNLS) domain or C-terminal putative autoregulatory domains (PADs) are absent (**Figure 6, Supplemental Figure S9**). This pNLS domain contains the conserved KGKAAL motif (KGKAAM in the case of OsWHY1), which is important not only for ssDNA binding (Krause et al., 2005; Grabowski et al., 2008), but also for RNA binding, as demonstrated in this study. In the crystal structure of StWHY1, Glu271 and Trp272 in the C-terminal PAD part are associated with Lys188 of the Whirly domain (Desveaux et al., 2005). This implies that the PAD region may be involved in the correct folding of the WHY1 protein. Deletion of the PAD region may prevent OsWHY1 from folding correctly and impair its RNA-binding activity. In addition, sequence alignment of twenty WHIRLY proteins from green algae to higher plants revealed that, apart from the highly conserved WHIRLY DNA-binding domain, both the N- and C-terminal sequences are quite divergent between taxa (Lin et al., 2024).

To further confirm the involvement of OsWHY1 in post-transcriptional processes of plastid genes, this study demonstrated that OsWHY1 directly targets specific sites in precursor RNAs of *rps14* and *rpl2* (**Figure 5, Supplemental Figure S8**). Interestingly, unlike certain PPR proteins which directly bind to RNA editing or splicing sites (Jiang et al., 2018; Xiao et al., 2018a; Xiao et al., 2018b; Lan et al., 2023), OsWHY1 either binds to the regions flanking *rps14-80* editing sites or the intronic region of the *rpl2* splicing site (**Figure 5, Supplemental Figure S8**), indicating that OsWHY1 likely binds to the periphery of the RNA modification sites of the targeted plastid transcripts. Considering the roles of WHY proteins in processing and stabilization of ssDNA intermediates during DNA replication, recombination and repair (Marechal et al., 2008; Marechal et al., 2009; Wang et al., 2021a), we speculated that OsWHY1 may play a stabilizing role in the RNA conformation, which likely facilitates the binding of other RNA editing factors or splicing factors to the correct RNA site for modification and processing. Recently, the cryo-electron microscopy structure of the tobacco PEP-PAP supercomplex revealed that the fifteen eukaryote-origin PAPs, binding to the periphery of the bacterial-origin PEP core, constitute four functionally distinct modules, in which the RNA module is formed by SMR-PPR proteins that may link transcription to RNA processing (Wu et al., 2024). However, WHY1 protein is not present in the PAP subunits of the RNA module. Thus, our findings provide a hint that OsWHY1-OsTRXz-OsMORFs complex may be involved in and assist the RNA processing by the RNA module of the PEP-PAP apoenzyme, but is not directly involved in the modification of target RNAs. The identification of more OsWHY1-interacting proteins in chloroplasts will help to refine the molecular regulatory mechanism of OsWHY1 in post-transcriptional modification of plastid genes.

### OsWHY1 recruiting OsTRXz-OsMORFs is involving in retrograde signaling pathway *via* alteration of the redox state and H_2_O_2_ homeotasis

Chloroplasts are the sites for operation of a series of photo-redox reactions leading to the production of ROS. The role of ROS as a signaling molecular implicated in stress responses and developmental senescence has long been established (Apel and Hirt, 2004; Miller et al., 2008). ROS are commonly toxic to cells and excessive accumulation of ROS in cells has detrimental effect on proteins, DNA and lipid, resulting in oxidative damage and cell death (Waszczak et al., 2018). Here, we found that the *oswhy1* mutants exhibit albino lethality and aberrant chloroplast development, which leads to photo-oxidative stress due to elevated production of ROS. (**Figure 1**; **Figure 7**). The transcription of genes related to ROS scavenging, such as *APXs* and *CATs*, was significantly decreased in *oswhy1* in contrast to the WT (**Figure 7C, 7D**). Plastid TRX and TRX-like proteins are known to play central roles in chloroplast protection against oxidative damage by regulating the redox balance (Cejudo et al., 2021; Yokochi et al., 2021). For example, OsTRXx and CDSP32 displayed thioredoxin activity in vitro and could reduce rice BAS1, a chloroplast 2-Cys PRX, which mediated hydrogen peroxide reduction (Perez-Ruiz et al., 2006). OsTRXm also exhibited reduction activity *in vitro* and regulated the activity of targeted 2-Cys PRX by reducing redox-active Cys residues. Knockdown of *ostrxm* resulted in pale-green leaves and defective chloroplast structure with increased H_2_O_2_ production (Chi et al., 2008). A cascade of TRX family genes, including *OsTRXm*, *OsTRXx* and *CDSP32*, were dramatically down-regulated in *oswhy1* compared with WT (**Figure 7D**), indicating an impaired redox balance caused by the loss of function of OsWHY1. The interplay between OsWHY1, OsTRXz and OsMORF8/9 suggests that OsWHY1/OsTRXz/OsMORFs complex may help modulate redox states to ensure H_2_O_2_ homeotasis and chloroplast stability. In fact, the increased ROS accumulation observed in *oswhy1, osmorf9* and *wp2/ostrxz* mutants validates the critical role of this regulatory module in chloroplast development (**Figure 9**).

Retrograde signals emanating from redox state of chloroplast often affect nuclear gene expression. Based on RT-qPCR and transcriptome analysis (**Figure 2**, **Figure 8**), we found that the upregulated gene sets in *oswhy1* albino seedlings were enriched for genes involved in various steps of chloroplast biogenesis and cytosolic translation, such as NEP-transcribed plastid housekeeping genes encoding the PEP apparatus and genes related to PEP activity, plastid tRNA/rRNA processing as well as plastid proteostasis, whereas downregulated genes were enriched for functions later in chloroplast development including PEP-mediated plastid gene transcription and PhANGs expression (**Figure 8**). These results point out that *OsWHY1* mutation activates plastid signals for chloroplast biogenesis, but disturbs retrograde signals for chloroplast development and photosynthetic differentiation probably by hampering plastid ribosome biogenesis and PEP activity. A possible explanation for this observation is that abnormal or immature chloroplast development gives rise to retrograde signals either promoting nuclear encoded plastid biogenesis program or repressing photosystem by downregulation of PhANGs as well as nuclear transcriptional activator of *PhANGs* (*OsGLK1*) (Nakamura et al., 2009; Leister and Kleine, 2016; Nagatoshi et al., 2016; Zubo et al., 2018). A recent finding also revealed that multiple plastid retrograde signals coordinately serve to distinct stages of chloroplast development by analyzing transcriptome dataset of non-photosynthetic maize mutants, where more than 170 plastid biogenesis-associated genes were up-regulated and most *PhANGs* were down-regulated (Kendrick et al., 2022), mirroring the transcriptome profile for our *oswhy1* mutant. The differential expression of nuclear genes in response to chloroplast dysfunction in *oswhy1* mutants underscores the importance of OsWHY1 in coordinating nuclear and plastid gene expression. It is consistence with the issue that WHY1 in Arabidopsis and barley have supposed to be involved in promoting photosynthetic gene expression under high light condition (Comadira et al., 2015; Huang et al., 2017). This coordination is crucial for the overall development and stress response of the plant, which is partially supported by the observation that overexpression of OsWHY1 appears to enhance chloroplast stability in leaves and increase photosynthetic efficiency under high light conditions (**Supplementary Figure S2**). It is very important for manipulation of high photosynthetic efficiency crop, with the help of OsWHY1 overexpression lines in the future.

In summary, OsWHY1 is integral to chloroplast development in rice through its RNA-binding activity and its role in recruiting OsTRXz and OsMORFs to ensure proper RNA modification. This understanding provides valuable insights into the molecular mechanisms of chloroplast development and expands our knowledge of the post-transcriptional regulatory functions of RNA-binding transcription factors on chloroplast genes.

## Materials and Methods

### Plant materials and growth conditions

The knockout mutants for *OsWHY1* in the ZH11 (*Oryza.Sativa L.ssp.japonica*) background were generated with the aid of Hangzhou Biogle Co., LTD. (Hangzhou, China) via CRISPR/Cas9 gene editing as described previously (Wang et al., 2015). The target gRNA sequence for *OsWHY1* was 5’-CCGCACCTCCTGCCTAGCCACAG-3’. The antisense lines were generated by transforming a *pTCK303* vector with a specific 137-bp cDNA fragment in the *OsWHY1* coding region into ZH11. For the overexpression lines, the full-length coding sequence of *OsWHY1* was cloned into the binary vector *pUN1301* under the control of the maize *UBQ* promoter, and the destination vector *pUN1301-OsWHY1,* in which the C-terminal of OsWHY1 was fused the GFP-tag, was subsequently transformed into the ZH11 using *Agrobacterium-*mediated transformation method (Jeon et al., 2000). The respective primers for cloning the coding sequences are listed in **Supplementary Table S2**. Rice seeds were germinated in wetted filter paper at 37 °C for two days in darkness and subsequently grown in an artificial climate chamber with a photoperiod of 12 h of light (∼250 μmol photons/ m^2^/s, 30 °C, 70 % humidity) and 12 h of darkness (25 °C, 70 % humidity). For the darkness and high light treatments, rice seedlings grown under normal light for 3 weeks were moved to either complete darkness or high-light conditions (∼360 μmol photons/ m^2^/s) for phenotype observation.

### Measurements of chlorophyll content and photosynthetic rate

To determine chlorophyll concentration, detached leaves were submerged in absolute ethanol solution to dissolve the pigments. After measuring the absorbance values of the solution by spectrophotometer (L3, INESA, China), total chlorophyll content was calculated based on the equations mentioned (Lichtenthaler and Wellburn, 1983) . Using a Pocket PEA Chlorophyll Fluorimeter (Hansatech Instruments, Norfolk, UK), the chlorophyll fluorescence of living leaves was measured as previously reported (Zheng et al., 2019). The average Fv/Fm of all primary leaves from five individual plants was calculated. An Imaging-PAM-Maxi (Walz, Effeltrich, Germany) was used to measure the chlorophyll fluorescence of living leaves and capture the image (Shao et al., 2008). Following a 30-minute adaptation period to total darkness, the maximal quantum yield of photosystem II (PS II; Fv/Fm) photochemistry and the minimum fluorescence at open PSII centers (Fo) were measured.

### Transmission electron microscopy analysis

The third fresh leaves from the *oswhy1* knockout mutants and the wild type ZH11 plants were collected and cut into small pieces. The leaf sections were prefixed in 2.5% glutaraldehyde (pH 7.4) for 4 h, rinsed with 0.1 M phosphate buffer (pH 7.2), postfixed overnight in 1% osmic acid at 4℃, dehydrated in a graded series of ethanol, and finally embedded in Epon-Araldite resin before ultrathin sectioning. Samples were stained with uranyl acetate and lead citrate, followed by observation under a Hitachi HT7700 transmission electron microscope.

### RNA isolation and gene expression analysis

The second fresh leaves from the *oswhy1* albino seedlings and its parallel wild type ZH11 plants were collected for RNA extraction. Total RNA was isolated from those leaf samples using TRIzol reagent according to the manufacturer’s instructions (Invitrogen) and treated with DNaseI. The mRNA was subjected to synthesize first-strand cDNA *via* the RevertAid First-Strand cDNA Synthesis Kit (Thermo Scientific™, Shanghai, China). Quantitative real time-PCR (qRT-PCR) was conducted to analyze the expression of genes using TransStart Green qPCR SuperMix Kit (TransGen Biotech, China) in the CFX96 machine (Bio-Rad Company, Hercules, CA, USA). The rice *OsACTIN* gene (*LOC_Os03g50885*) was selected as the reference gene. The 2^-ΔCt^ method was used to normalize expression levels of tested genes as described (Schmittgen and Livak, 2008). The experiment was performed three biological replicates. The primer sequences used for the expression analysis were listed in **Supplementary Table S2**.

### Transcriptome analysis

Total RNA was isolated from 7-day-old seedlings of *oswhy1* albino mutants and wild type. RNA purity and quality was determined using a Nanodrop spectrophotometer and Agilent 2100 analyzer. Direct RNA-sequencing libraries were generated and sequenced based on Oxford Nanopore Technologies (ONT) third-generation sequencing platform (Garalde et al., 2018) by Beijing BioMarker Technologies (Beijing, China). Clean reads were mapped to the TIGR7 genome using TopHat 2.0.11 (Trapnell et al., 2009). The gene transcript levels were analyzed by the CPM (counts per million)method (Zhou et al., 2014). Differentially expressed genes (DEGs) were determined using DESeq2 according to a threshold an absolute value |log2_ratio|≥1 and FDR≤0.01. All of the DEGs were used for the GO, KEGG and functional annotation analyses (Ashburner et al., 2000; Kanehisa et al., 2004). Two independent biological replicates were performed for the RNA-seq.

### Plastid RNA editing/splicing analysis

Using the cDNAs obtained from the wild type and *oswhy1* albino seedlings as templates, the PCR products of the corresponding plastid genes were amplified with their respective primers around the editing sites (**Supplementary Table S2**) and sequenced directly. The C to T editing levels of each site were measured according to the sequencing profiles as mentioned previously (Tian et al., 2019). For RNA splicing analysis, the chloroplast genes with at least one intron were selected and amplified with primers flanking the introns (**Supplementary Table S2**). The experiments were conducted three biological replicates.

### Subcellular localization analysis and confocal microscopy

With the aid of the Gateway® cloning recombination technology, the full-length cDNAs of *OsWHY1* (*LOC_Os06g05350*), *OsMORF1* (*LOC_Os11g11020*), *OsMORF8* (*LOC_Os09g33480*), *OsMORF9* (*LOC_Os08g04450*) and *OsTRXz* (*LOC_Os08g29110*) were cloned into the entry vector *pDONR201* and, subsequently, recombined into the expression vectors *p2GWF7* (C-terminal GFP fusion) and *p2GWR7* (C-terminal RFP fusion) under the control of *Cauliflower Mosaic virus* (CaMV35S) promoter as described previously (Zheng et al., 2019). To construct the expression vector of OsWHY1 deletion variant, the *OsWHY1* gene without the region encoding the putative chloroplast transit peptide (51-272aa) was amplified by PCR, cloned into *pDONR201*, followed by recombination into *p2GWF7*. Primers used for these vector constructions are listed in **Supplementary Table S2**. All constructs were confirmed by sequencing. The expression vector carrying *p35S:OsWRKY44-RFP* was described by Habiba et al. (2021).

For subcellular localization observation, the corresponding expression constructs were separately transformed into rice protoplasts as previously described (Zheng et al., 2019). For the co-localization assay, the expression vector of OsWHY1-GFP or OsWHY1 51-272aa-GFP was co-transfected with OsWRKY44-RFP or OsMORFs-RFP into rice protoplasts according to the description (Habiba et al., 2021) and incubated overnight at room temperature for protein expression. All microscopic observations were monitored using a Leica TCS SP8 confocal laser scanning microscope as before described (Zheng et al., 2019). Excitation/emission wavelengths were 488nm/505-535nm for GFP, 561nm/575-605nm for RFP and 633nm/650–710 nm for chloroplast auto-fluorescence. Image analysis was processed with Leica TCS SP8 software, Adobe Photoshop and Adobe Illustrator.

### Yeast two-hybrid (Y2H) analysis

After amplification with gene-specific primers (**Supplementary Table S2**), the full-length cDNAs of *OsTRXz*, *OsPGL1 (LOC_Os12g06650), OsPPR57940 (LOC_Os04g57940), OsPPR42120 (LOC_Os12g42120)* and *OsPPR24930 (LOC_Os05g24930)* were integrated into *EcoRI/SacI* sites of the *pGADT7* vector, and *OsWHY1* coding sequence was cloned into *EcoRI/SalI* sites of the pGBKT7 vector *via* the ClonExpress® II recombinant Cloning Kit (Vazyme, C112-01). The constructs of *pGADT7-OsMORF1*, *pGADT7-OsMORF2*, *pGADT7-OsMORF3*, *pGADT7-OsMORF8*, *pGADT7-OsMORF9*, *pGADT7-OsWSP1* were kindly provided by Prof. Dr. Hu Jun of Wuhan University (Xiao et al., 2018a). Different combinations of bait and prey constructs were subsequently co-transformed into yeast strain AH109 for the Y2H assays as described previously (Huang et al., 2022).

### Bimolecular fluorescence complementation (BiFC) analysis

The plasmids *pRTVcVN* and *pRTVcVC* were used for BiFC assay (He et al., 2018a). Full-length coding sequences of *OsWHY1* and candidate proteins without the stop codon were inserted into *pRTVcVC* (C-terminal fused with C-truncated Venus fluorescent protein) and *pRTVcVN* (C-terminal fused with N-truncated Venus fluorescent protein), respectively, using the ClonExpress® II recombinant Cloning Kit (Vazyme, C112- 01). The primers used to generate BiFC constructs are listed in **Supplementary Table S2**. For transient expression, different combinations of BiFC constructs were co-transformed into rice protoplasts obtained from 2-week-old seedlings as described previously (Zheng et al., 2019). After incubation 16h, the transformed protoplasts were examined *via* a Leica TCS SP8 confocal laser scanning microscope (Zheng et al., 2019). The excitation and detection wavelengths for Venus was 514 nm for excitation and 520-570 nm for detection.

### Co-immunoprecipitation (Co-IP) and immunoblotting assays

The full-length coding region of *OsWHY1*, with a C-terminal HA tag, was amplified using gene-specific primers (Supplementary Table S2) and cloned into *pDONR201*. This construct was subsequently recombined into the expression vector *p2GW7* (Life Technologies, Darmstadt, Germany) using Gateway® cloning technology. The plasmid *p2HAGW7* (N-terminal HA-tag) served as an empty vector control. Constructs expressing the corresponding HA-tagged and GFP-tagged proteins were co-transformed into rice protoplasts. After a 16-hour incubation, the transfected protoplasts were harvested for Co-IP and immunoblotting assays, following the protocol described previously (Zheng et al., 2018). The HA-tagged proteins were immunoprecipitated using anti-HA antibody-coupled beads (anti-HA affinity matrix, Roche), while the GFP-tagged proteins were immunoprecipitated using anti-GFP affinity beads (SA070001; Smart-Lifesciences, Changzhou, China). The immunoprecipitated proteins were then separated via 13.5% SDS-PAGE, blotted, and analyzed by immunoblotting using anti-GFP (TransGen, HT801, 1:1,000 dilution) and Anti-HA-Peroxidase (Roche, 3F10, 1:500 dilution) antibodies.

### Protein extraction and western blot analysis

Total proteins were extracted from young leaves of 10-day-old wild-type and oswhy1 mutant seedlings. 100 mg leaves were collected and grounded to fine powder in liquid nitrogen. After the addition of 100 μl of ice-cold extraction buffer (50mM HEPES pH 7.4, 150 mM NaCl, 0.1 % Trition X-100, 1 mM EDTA, 1 mM DTT, 1 x phosphatase inhibitor cocktail and 1 x protease inhibitor cocktail) and incubation on ice for 30 minutes, the plant tissue was further homogenized by vortex mixing. The soluble proteins were purified from the mixture by centrifugation at 4 °C, 15000 g for 20 minutes and denatured in protein loading buffer at 95 °C for 10 min. The proteins were loaded onto a 13.5% SDS-polyacrylamid gel and separated by electrophoresis (SDS-PAGE). PageRuler Prestained protein ladder (Fermentas) was used as a molecular weight marker. Proteins were transferred to nitrocellulose membranes (Hybond–ECL, Amersham) and stained with 0.1% Ponceau S to visualize sample loading. The relevant antibodies were obtained from BPI (http://www.proteomics.org.cn/).

### RNA electrophoresis mobility shift assays (REMSAs)

The *pCold-GST* vector was used for the production of recombinant protein in *Escherichia coli* (Hayashi and Kojima, 2008). The corresponding cDNA fragments of OsWHY1 were amplified with specific primers and cloned into the *pCold-GST* to create the constructs encoding GST- OsWHY1 51-272aa, GST-OsWHY1 96-272aa, GST-OsWHY1 128-272aa, GST-OsWHY1 51-235aa or GST-OsWHY1 97-235aa fusion proteins. These recombinant proteins were expressed at 15°C with 0.5 mM isopropyl-β-D-thiogalactopyranoside (IPTG) for 20 h and purified by Glutathione Magarose Beads (SM002005, Smart-Lifesciences, Changzhou, China). The RNA probes (*rps14-p1*, *rps14-p2, rps14-p3, rpl2-p1, rpl2-p2* and *rpl2-p3*) were synthesized and labeled with Cy5 at the 5ʹ end by GenScript (Nanjing, China). For RNA electrophoresis mobility shift assays (REMSAs), the recombinant protein (1 μg) was incubated with the Cy5-labelled RNA probe (1 μM) in a 20 μl reaction mixture including 4 μl of 5×binding buffer (50 mM Tris-HCl pH7.5, 10 mM NaCl, 200 mM KCl, 5 mM MgCl2, 10 units RNasin, 10 mM EDTA pH8.0, 5 mM dithiothreitol and 25 mg/L BSA). The mixture was incubated at 25 °C for 1 h followed by separation in 6% native polyacrylamide gel electrophoresis (PAGE) in 0.5× Tris-borate-EDTA (TBE) buffer. Electrophoresis was performed on a cooled support at 110 V for 60 minutes. The fluorescence measurement of the polyacrylamide gel was detected on a G:BOX Chemi XT4 gel system at 635 nm for excitation and 700 nm for emission. The images of the shifts were inverted to black-and-white. For the competitive REMSAs, a gradually increasing concentration of unlabelled probe was added to the binding reaction according to the procedure described above.

### NBT, DAB staining and H_2_O_2_ content measurement

Qualitative analysis of superoxide free radicals and hydrogen peroxide (H_2_O_2_) were carried out using Nitro blue Tetrazolium (NBT) and 3, 3′-diaminobenzidine (DAB) staining, respectively, following methods described previously (Huang et al., 2019; Lin et al., 2020) with minor modifications. The second-fully expended leaves of 7-day-old rice seedlings were harvested and immersed in 0.5 mg/mL NBT solution (50 mM sodium phosphate buffer, pH 7.5) or 1.0 mg/mL DAB solution (50 mM Tris-HCl, pH 5.0). After vacuum infiltration, the samples were incubated at 28°C for 8 h in darkness. After that, the stained samples were cleared in boiling ethanol (95%) for 20 minutes and incubated further for 24 h at room temperature to remove the chlorophyll. Imaging was conducted using an Epson Perfection V600 Photo scanner. For quantitative measurement of H_2_O_2_ production, H_2_O_2_ was extracted from leaves by cold acetone (Ruan et al., 2011) and quantified using the Hydrogen Peroxide Content Assay Kit (Sangon Biotech Co, Ltd., D799774).

### Statistical analysis

All experiments involving measurements, quantifications and imaging were performed at least in triplicate with similar results. The Student’s t-test was used to determine significant difference compared to the wild type control (* P < 0.05, **P < 0.01). Data plotting and statistical tests were carried out using the GraphPad Prism software version 8 (GraphPad Software, San Diego, CA, USA). The values presented in the figures are means ±SD.

### Accession numbers

Sequence data from this article can be found in the GenBank database under the following accession numbers: OsWHY1, LOC_Os06g05350; OsWHY2, LOC_Os02g06370; OsTRX z, LOC_Os08g29110; OsMORF1, LOC_Os11g11020; OsMORF2, LOC_Os06g02600; OsMORF3, LOC_Os03g38490; OsMORF8, LOC_Os09g33480; OsMORF9, LOC_Os08g04450; OsWSP1, LOC_Os04g51280; OsPGL1, LOC_Os12g06650; OsPPR57940, LOC_Os04g57940; OsPPR42120, LOC_Os12g42120; OsPPR24930, LOC_Os05g24930; OsACTIN, LOC_Os03g50885; HSP, LOC_Os09g30418; RPL2, LOC_Osp1g00770; RPS14, LOC_Osp1g00320; RPS4, LOC_Osp1g00360; rpoA, LOC_Osp1g00660; rbcL, LOC_Osp1g00420; PsbD, LOC_Osp1g00170.

## Supplementary Information

Supplementary Table S1. Differentially expressed genes (DEGs) were counted in *oswhy1* vs WT Supplementary Table S2. Sequence of the primers used in this study

Supplementary Table S3. Potential OsWHY1-interacting proteins identified by LC-MS/MS analysis.

Supplementary Figure S1. Subcellular localization of OsWHY1.

Supplementary Figure S2. Phenotypic Analysis of OE-OsWHY1-GFP Overexpression Lines. Supplementary Figure S3. Differentially expressed genes (DEGs) involved in photosynthesis in the *oswhy1* mutants.

Supplementary Figure S4. OsWHY1 interacts with OsTRXz. Supplementary Figure S5. OsWHY1 interacts with OsMORF proteins.

Supplemental Figure S6. RNA editing analysis of chloroplast transcripts in the WT and *oswhy1* mutant plants.

Supplemental Figure S7. Splicing analysis of chloroplast transcripts with introns in rice WT and the two oswhy1 mutant plants.

Supplemental Figure S8. REMSA analysis of OsWHY1 protein. Supplemental Figure S9. REMSA analysis of OsWHY1 deletion mutants. Supplementary Figure S10. Phenotype of Oswhy1 RNAi rice plants.

## Acknowledgements

We are grateful to Prof. Dr. Hong-Wei Xue (School of Agriculture and Biology, Shanghai Jiao Tong University) and Dr. Yanxiang Zhang for their technical assistance to generate overexpression transgenic lines and RNAi mutant lines of *OsWHY1*. We thank Prof. Dr. Jun Hu (College of Life Sciences, Wuhan University) for providing us with the Y2H vertors expressing OsMORF1, OsMORF2, OsMORF3, OsMORF8, OsMORF9 or OsWSP1. We are also grateful to Prof. Dr. Jianmin Wan (State Key Laboratory of Crop Genetics and Germplasm Enhancement, Nanjing Agricultural University) for gifting us seeds of the *wp2* mutant.

## Author Contributions

X.Z.Z and Y.M conceived and designed the research. X.Z.Z, Q.Z.L, Y.L.L and J.X.X performed experiments and collected the data. X.Z.Z and Y.M draft and revised the manuscript. Other authors assisted in experiments, data analysis and discussed the results. All authors read and approved this manuscript.

